# Intellectual disability-associated factor Zbtb11 cooperates with NRF-2/GABP to control mitochondrial function

**DOI:** 10.1101/2019.12.13.875708

**Authors:** Brooke C. Wilson, Lena Boehme, Ambra Annibali, Alan Hodgkinson, Thomas S. Carroll, Rebecca J. Oakey, Vlad C. Seitan

**Author notes:** These authors contributed equally (Brooke C. Wilson, Lena Boehme, Ambra Annibali).

## Abstract

Zbtb11 is a conserved transcription factor mutated in families with hereditary intellectual disability. Its precise molecular and cellular functions are currently unknown, precluding our understanding of the aetiology of this disease. Using a combination of functional genomics, genetic and biochemical approaches here we show that Zbtb11 plays essential roles in maintaining the homeostasis of mitochondrial function. Mechanistically, we find Zbtb11 facilitates the recruitment of Nuclear Respiratory Factor 2 (NRF-2) to its target promoters, activating a subset of nuclear genes with roles in the biogenesis of respiratory complex I and the mitoribosome. Genetic inactivation of *Zbtb11* resulted in a severe complex I assembly defect, impaired mitochondrial respiration, mitochondrial depolarisation, and ultimately proliferation arrest and cell death. Experimental modelling of the pathogenic human mutations showed these have a destabilising effect on the protein, resulting in reduced Zbtb11 dosage, down-regulation of its target genes, and impaired complex I biogenesis. Our study establishes Zbtb11 as a novel essential mitochondrial regulator, improves our understanding of the transcriptional mechanisms of nuclear control over mitochondria, and provides a rationale for the aetiology of Zbtb11-associated intellectual disability.

## Introduction

The biochemical energy contained within nutrients is captured through the process of cellular respiration, which culminates in the production of ATP through oxidative phosphorylation (OXPHOS) in mitochondria (Cooper & Lehninger, 1957). OXPHOS is fuelled by a chain of redox reactions in which electrons are transferred along an electron transport chain (ETC) comprised of 4 complexes (I-IV) localised to the mitochondrial inner membrane (Dudkina *et al*, 2008). Three of the ETC complexes (I, III and IV) couple substrate oxidation with proton extrusion from the mitochondrial matrix into the intermembrane space, thus creating an electrochemical gradient across the mitochondrial inner membrane which forces protons back into the matrix mainly through complex V (ATP synthase), driving the synthesis of ATP (Schultz & Chan, 2001). While mitochondria are ubiquitous organelles, tissues with high energy demands, such as brain and muscle, are particularly reliant on the activity of the ETC, and these are usually disproportionally affected in mitochondrial diseases. Nevertheless, because the activity of the ETC is also vital for regenerating the cellular pool of redox cofactors that catalyse a number of other metabolic reactions (Birsoy *et al*, 2015; Sullivan *et al*, 2015), it is becoming increasingly evident that a functional ETC is essential for all proliferating cells, irrespective of their ATP consumption (Yao *et al*, 2019).

Mitochondria possess their own genome (mtDNA) and transcription machinery, as well as a specialised organellar ribosome (the mitoribosome). While the mtDNA encodes a small number of core subunits of the OXPHOS complexes, the majority of the mitochondrial proteome is encoded in the nuclear genome (nDNA). Mitochondrial biogenesis and function are therefore controlled to a significant extent through transcriptional regulation of nuclear genes. A number or transcription factors have been implicated in the regulation of nuclear-encoded mitochondrial genes, some of which are essential while others have modulatory roles important mainly in specific cell types or in conditions of increased energy demand (Fernandez-Marcos & Auwerx, 2011; Scarpulla *et al*, 2012). The nuclear respiratory factors (NRFs) NRF-1 and NRF-2 (also known as GABP) are essential transcriptional activators, and the most prominent regulators of nuclear-encoded mitochondrial genes, driving the expression of the majority of nuclear-encoded OXPHOS subunits, mitoribosomal proteins, as well as mtDNA replication and transcription factors (Quirós *et al*, 2016). NRF-1 and NRF-2/GABP play overlapping but non-redundant roles in activating these genes, and they are found associated with different DNA sequence motifs. It is not currently completely understood how the activity of NRFs is regulated, how their recruitment to target genes is controlled, or whether they engage all their genomic targets in the same way or through locus-specific mechanisms.

Zinc finger and BTB domain-containing (ZBTB) proteins are a family of structurally related transcription factors, which function by binding DNA through C-terminal zinc finger motifs and recruiting co-factors via protein-protein interactions mediated by an N-terminal BTB domain (Lee & Maeda, 2012). Zbtb11 is a poorly characterised ubiquitously expressed ZBTB protein that is conserved in vertebrates. Recent genetic studies of consanguineous families with recessive intellectual disability (ID) identified homozygous missense mutations in human *ZBTB11* as causal variants (Harripaul *et al*, 2017; Fattahi *et al*, 2018). Two pathogenic mutations were identified, both of which are predicted to disrupt individual Zbtb11 zinc finger motifs. Affected patients display morphological defects in the brain, including ventriculomegaly and cerebellar atrophy, and also show neuromuscular defects such as ataxia and facial hypotonia (Fattahi *et al*, 2018, 11). The molecular and cellular functions of Zbtb11 have hitherto remained unknown, so the aetiology of this disease is not currently understood.

Here we investigate the cellular and molecular functions of Zbtb11 and find that it is a key regulator of mitochondrial function. We show that Zbtb11 controls the locus-specific recruitment of NRF-2/GABP, but not NRF-1, to activate a subset of nuclear-encoded genes with roles in the biogenesis of OXPHOS complex I and the mitoribosome. Genetic inactivation of Zbtb11 leads to a severe complex I assembly defect and loss of complex I activity, reduced respiration, mitochondrial depolarisation, and consequently to proliferation arrest and cell death. Zbtb11 therefore cooperates with NRF-2/GABP to maintain the homeostasis of mitochondrial function. We provide evidence that mutations associated with hereditary intellectual disability disrupt this function by destabilising the Zbtb11 protein, leading to its reduced dosage, down-regulation of its target genes and impaired complex I biogenesis. Our study establishes Zbtb11 as an essential regulator of mitochondrial function, reveals a previously unanticipated mechanism of locus-specific regulation of NRF-2/GABP activity, and indicates that intellectual disability associated with mutations in *ZBTB11* may be at least in part the manifestation of a mitochondrial disease.

## Results

### Zbtb11 is enriched at promoters of nuclear-encoded mitochondrial genes

We initially set out to determine the potential functions of Zbtb11 by identifying the exact locations where it binds in the genome, and to this end we carried out chromatin immunoprecipitation coupled with high-throughput sequencing (ChIP-seq) experiments. Due to lack of existing validated anti-Zbtb11 antibodies, we used CRISPR/Cas9 to ‘knock-in’ a 3xFLAG tag into the *Zbtb11* locus of the E14 mouse ESC line (Doetschman *et al*, 1987) (Fig. 1a), and isolated a homozygous line in which all endogenous Zbtb11 protein is N-terminally tagged with 3xFLAG (FLAG-Zbtb11) (Fig. 1b and Supplementary Fig.1a). We then performed anti-FLAG ChIP-seq in *Zbtb11*^FLAG/FLAG^ E14 cells and control E14 WT cells, identifying 9,350 FLAG-Zbtb11 consensus peaks from 3 biological replicates (Irreproducibility Discovery Rate (IDR) < 0.02, see Methods), from which we removed 219 peaks that that were also identified by FLAG ChIP-seq in control E14 WT cells.

**Fig. 1.**
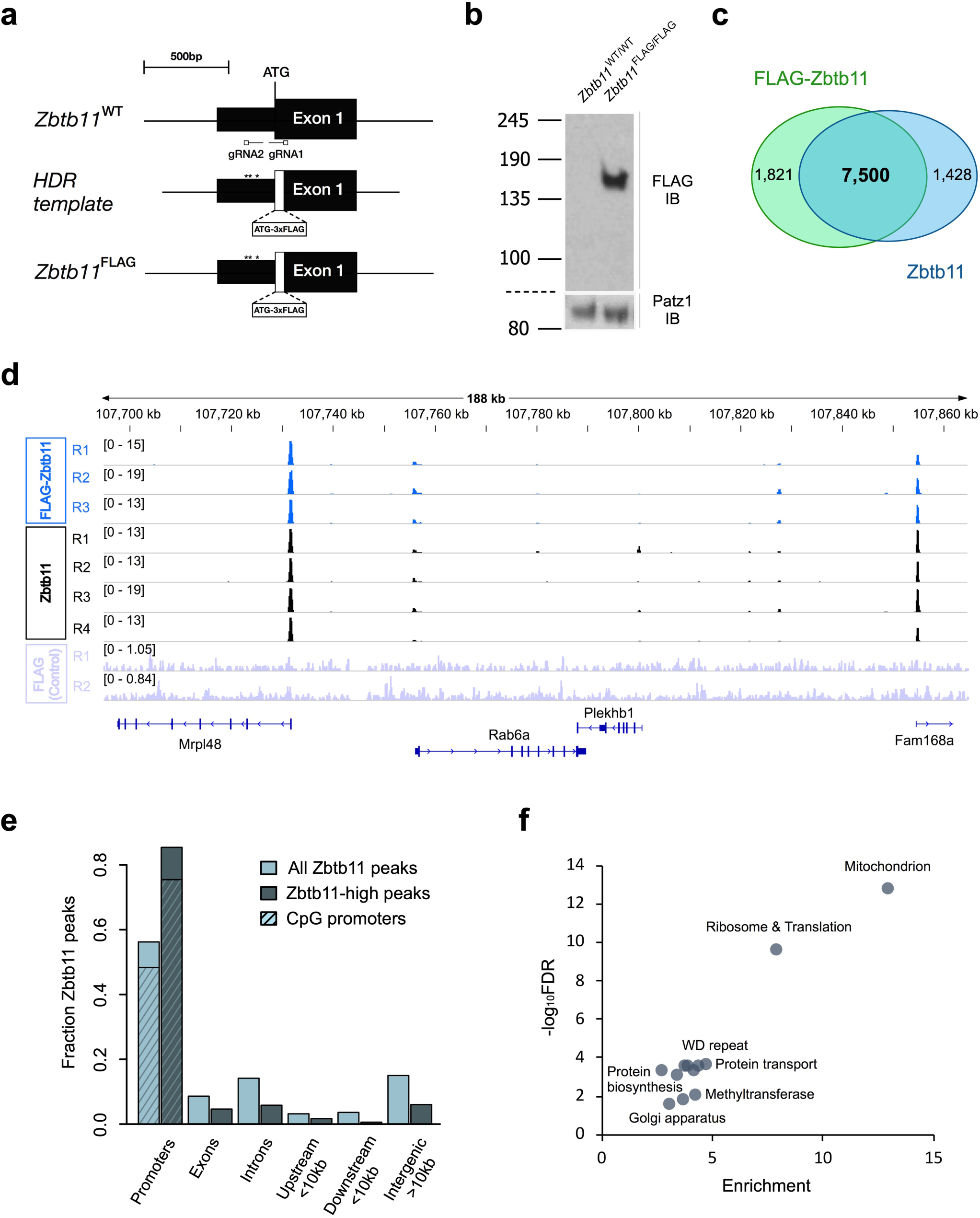
Zbtb11 is enriched at promoters of nuclear-encoded mitochondrial genes. (a) CRISPR/Cas9 ’knock-in’ approach used to generate the *Zbtb11*^FLAG^ allele. Mouse E14 ESCs were co-transfected with two gRNAs targeting the 5’ end of the coding region in the *Zbtb11* gene, the Cas9 nickase, and a donor template for homology-directed repair (HDR) which contained the 3xFLAG tag immediately downstream of, and in the same open reading frame (ORF) with the translation initiating methionine. (*) - approximate position of poorly conserved non-coding single nucleotides that were changed in the donor template to avoid targeting by gRNA2. The sequence recognised by gRNA1 is disrupted in the HDR template by the insertion of the 3xFLAG sequence. (b) Anti-FLAG immunoblot of whole cell lysates from a *Zbtb11*^FLAG/FLAG^ homozygous cell line, and the parental line E14. Patz1 was used as loading control. (c) Genome-wide overlap between ChIP-seq peaks obtained by performing FLAG ChIP in *Zbtb11*^FLAG/FLAG^ E14 cells and ChIP with anti-Zbtb11 antibodies in the parental line E14 (see also Supplementary Fig.1c). (d) ChIP-seq signal at an example Zbtb11 target locus (*Mrpl48*). FLAG ChIP in E14 WT (FLAG control) is shown as background control (see also Supplementary Fig.1c and S1D). (e) Genome-wide distribution of Zbtb11 peaks with respect to genes. The distribution of the entire Zbtb11 ChIP-seq dataset is shown alongside that of Zbtb11-high peaks. (f) Annotation terms associated (FDR < 0.05) with genes that have Zbtb11-high peaks present at their promoters. Plot shows enrichment values vs significance (-log_10_FDR) when compared in DAVID against all genes in the mouse genome.

We used this reliable set of FLAG-Zbtb11 binding sites to validate an anti-Zbtb11 antibody which, although not suitable for immunoblotting, reproducibly showed strong ChIP enrichment at the same sites as FLAG-Zbtb11 (Supplementary Fig.1b). ChIP-seq with this Zbtb11 antibody in E14 WT cells detected 8,957 Zbtb11 peaks (IDR < 0.02), of which 7,500 overlapped with the FLAG-Zbtb11 peaks in *Zbtb11*^FLAG/FLAG^ E14 cells (Fig. 1c). The ChIP-seq signal was very well correlated between Zbtb11 and FLAG-Zbtb11 data sets (Fig. 1d and Supplementary Fig.1c and 1d). We therefore generated a high-confidence Zbtb11 peak set comprised of the 7,500 peaks common to both Zbtb11 and FLAG-Zbtb11, which we subsequently used in the rest of the analyses in this study.

We found Zbtb11 binds predominantly (56%) at gene promoters, the vast majority (85.6 %) of which are CpG island promoters (Fig. 1e) - a promoter type mainly associated with ubiquitously expressed housekeeping genes (Ramsköld *et al*, 2009; Vavouri & Lehner, 2012). Analysing the distribution of Zbtb11 peak signal values revealed that the top 858 peaks are significantly stronger than the rest of the peaks in the data set, to the extent that they are outliers in the distribution (values are higher than 1.5 times the inter-quartile range over the third quartile) (Supplementary Fig.1f). These Zbtb11-high peaks represent the top 10% of the peak set, and their signal is overall 8.7 times stronger than the rest of the peaks. Zbtb11-high peaks show increased association with promoters compared to the entire Zbtb11 peak data set (86.2% vs 56%) (Fig. 1e).

To identify pathways and cellular processes potentially controlled by Zbtb11, we carried out functional enrichment analyses using genes with the most prominent Zbtb11 binding sites (the 858 Zbtb11-high peaks). The most highly enriched annotation term cluster was related to mitochondria (13-fold enrichment), 15% of tested genes being included in this category (Fig. 1f). This was followed at some distance by ribosome and translation (7.8-fold enrichment), however 52% of genes in this category function in mitochondrial translation and were also listed in the mitochondria annotation cluster. We did not detect any enrichment of terms related to development or ESC differentiation. Zbtb11 therefore seems to mainly associate with promoters of a subset of housekeeping genes, among which genes with mitochondrial functions are highly enriched.

### Zbtb11 is essential for proliferation and cell viability

To establish an experimental system that would allow us to interrogate the functions of Zbtb11 we first attempted to generate constitutive *Zbtb11* KO ESCs by targeting exon 3 using CRISPR/Cas9-mediated genome editing. However, although we were able to efficiently isolate and expand heterozygous WT/KO clones, we were never able to isolate homozygous *Zbtb11* KO lines, which suggested that *Zbtb11* is essential for cell viability. We therefore took a different approach, generating an inducible *Zbtb11* KO ESC line. Using CRISPR/Cas9-mediated ‘knock-in’, we flanked exon 3 of *Zbtb11* with loxP sites, and also inserted the ERt2-Cre transgene (Indra *et al*, 1999) in the *Rosa26* (*R26*) locus (Zambrowicz *et al*, 1997) (Fig. 2a and Supplementary Fig.2a). In these cells ERt2-Cre can be induced by addition of 4-hydroxytamoxifen (4OHT) to delete *Zbtb11* exon 3, which is predicted to induce a frame shift and truncate the Zbtb11 protein to an 185 amino acid N-terminal fragment that lacks the BTB domain and all of the zinc fingers motifs.

**Fig. 2.**
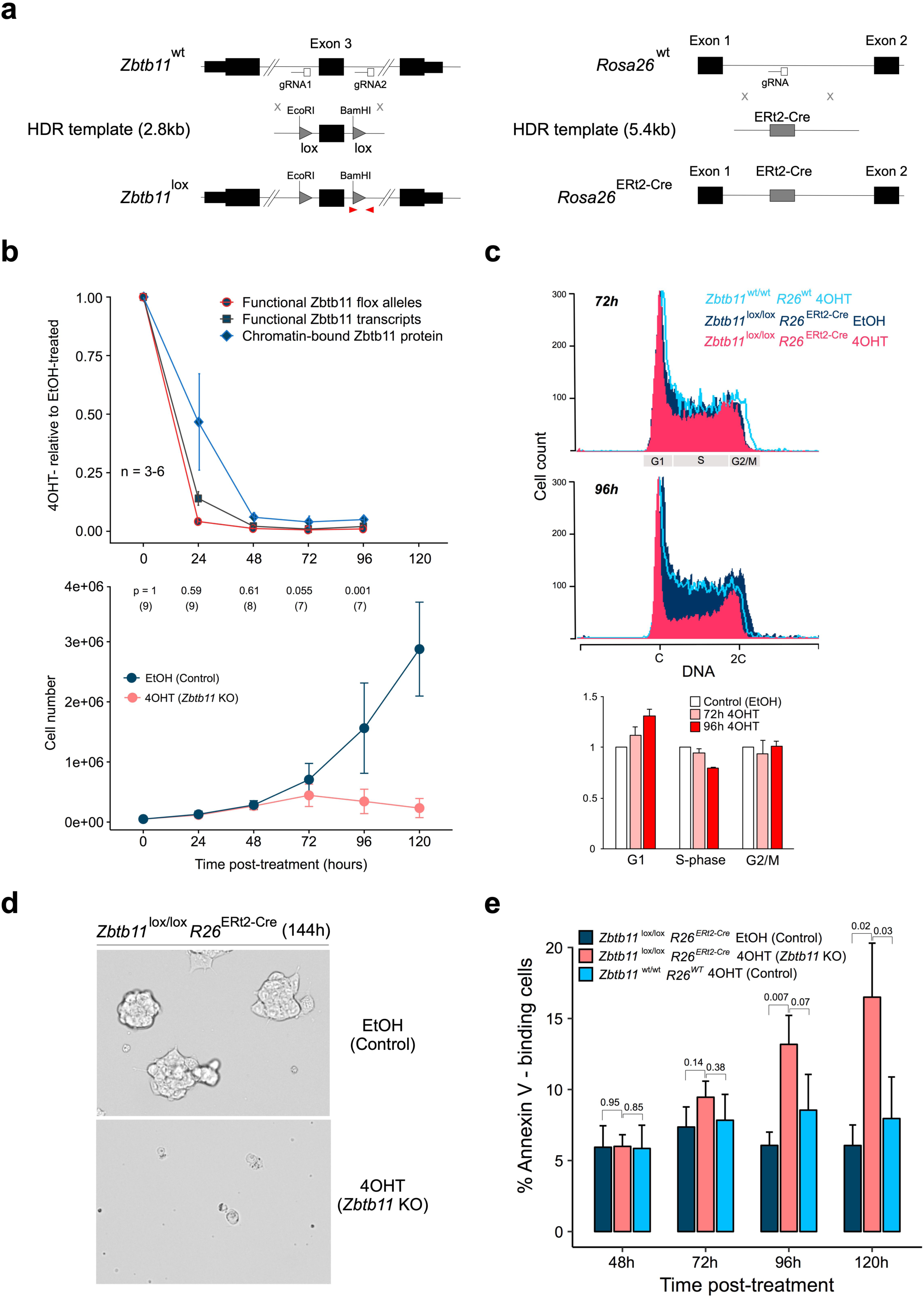
Zbtb11 is essential for proliferation and cell viability. (a) CRISPR ’knock-in’ strategies used to generate the conditional *Zbtb11* KO allele (*Zbtb11*^lox^) (left), and to insert the ERt2-Cre transgene in the *Rosa26* locus (right). The loxP sites (lox) flanking *Zbtb11* exon 3 include ectopic restriction sites (see Supplementary Fig.2a). Red arrow heads mark qPCR primers used to measure the deletion efficiency once ERt2-Cre is activated - recombination between the two loxP sites leads to excision of the left primer binding site along with exon 3 so PCR products are no longer generated. (b) Time-course experiment describing the dynamics of Zbtb11 inactivation in our inducible *Zbtb11* KO ESC line (*Zbtb11*^lox/lox^ *Rosa26*^ERt2-Cre^). Upper panel - decay of functional *Zbtb11* alleles, *Zbtb11* transcripts and protein in 4OHT-treated cells relative to their EtOH-treated control cells following KO induction. Lower panel - growth curves (mean cell counts ± SD), with p-values (paired t-test) above and number of replicates in brackets. (c) Propidium iodide stain for cell cycle analysis of control (*Zbtb11*^lox/lox^ EtOH) and *Zbtb11* KO (*Zbtb11*^lox/lox^ 4OHT) cells. 4OHT-treated *Zbtb11*^wt/wt^ cells were included to control for potential non-specific effects of 4OHT. Top two panels - representative flow cytometry histograms showing the distribution of DNA content in the sampled cell populations. G1, S or G2/M phases of the cell cycle are indicated on the horizontal axis. Bottom panel - mean and SD of the fractions of 4OHT-treated cells present in each cell cycle phase (normalised to EtOH-treated controls, n = 3). (d) Bright field microscopy images of 144 hours *Zbtb11* KO and control cells, illustrating the collapse of the *Zbtb11* KO culture at this time point. (e) Quantification of cell death after *Zbtb11* KO induction with FITC-coupled Annexin V. Bars show fraction of Annexin V-positive cells (mean and SD of 3-5 biological replicates) (see also Supplementary Fig.2d). P-values (two-tailed t-test) are shown above.

Treatment of *Zbtb11^lox/lox^ R26^ERt2-Cre^* cells with 4OHT, but not with the carrier ethanol (EtOH), resulted in efficient deletion of *Zbtb11* exon 3 and depletion of functional Zbtb11 mRNA transcripts, which was essentially complete 24 hours after 4OHT treatment (Fig. 2b). ChIP-qPCR experiments with anti-Zbtb11 antibodies showed the signal was decreased 24 hours after treatment and became virtually undetectable from 48 hours post-treatment onwards (Fig. 2b and Supplementary Fig.2b).

This further validated the specificity of the anti-Zbtb11 ChIP antibody, and showed that Zbtb11 is completely depleted at the protein level 48 hours after *Zbtb11* KO is induced. Acute depletion of Zbtb11 transcripts and protein is maintained at 72 and 96 hours post-KO induction (Fig. 2b), indicating there is no selection for cells that may escape deletion and have a growth advantage. These results establish our inducible KO cell line as a robust experimental system that allows rapid and acute depletion of Zbtb11 to study its cellular functions.

*Zbtb11^lox/lox^ R26^ERt2-Cre^* cells treated with EtOH continued to grow exponentially as long as they were regularly passaged. By contrast, the proliferation of 4OHT-treated cells slowed down after Zbtb11 was completely depleted from the chromatin, being completely arrested 96 hours after treatment (Fig. 2b). Cell cycle analyses by flow cytometry revealed that Zbtb11 depletion resulted in a reduction of cells in S-phase and a relative increase of cells in G1, while the proportion of cells in G2/M did not appear to change (Fig. 2c). This indicates that the slow-down in proliferation caused by Zbtb11 depletion is mainly underpinned by cell cycle arrest in G1. Concomitant with proliferation arrest, cell death was also noticeable in 4OHT-treated cultures starting at day 4 (96h), becoming extensive a day later (120h) so that very few cells were left in culture beyond 5 days post-treatment (Fig. 2d). Flow cytometry measurements of cleaved Caspase-3 failed to detect any increase in apoptosis rates (Supplementary Fig.2c). However, the increase in cell death was measurable by Annexin V-mediated detection of phosphatidylserine externalisation (Fig. 2e and Supplementary Fig.2d), which was significant from 96 hours post-4OHT treatment onwards. No significant changes in proliferation or cell death were discernible in 4OHT-treated *Zbtb11^wt/wt^ R26^wt^* cells (Fig. 2c and 2e), confirming that the observed effects were not artefacts of 4OHT treatment. These results establish that Zbtb11 is essential for proliferation and cell viability, and when its functions are blocked cells undergo cell cycle arrest in G1 and subsequently suffer Caspase-independent cell death.

### Zbtb11 supports the expression of a subset of nuclear-encoded mitochondrial proteins

To determine what processes are disrupted by the depletion of Zbtb11, we carried out whole transcriptome analyses by directional RNA-sequencing (RNA-seq). We compared *Zbtb11^lox/lox^ R26^ERt2-Cre^* cells treated with either 4OHT (*Zbtb11* KO) or EtOH (control). To control for non-specific effects of 4OHT we also analysed *Zbtb11^wt/wt^ R26^wt^* cells treated with 4OHT. We analysed cells treated for 48 hours because at this time point Zbtb11 protein is completely depleted in 4OHT-treated cells (see Fig. 2b and Supplementary Fig.2b), but there are no detectable changes in proliferation and cell survival (see Fig. 2c and 2e). We therefore expected to detect mainly transcriptional changes at genes directly regulated by Zbtb11.

When comparing 4OHT- and EtOH-treated *Zbtb11^lox/lox^ R26^ERt2-Cre^* cells, we found 154 differentially expressed (DE) genes (FDR < 0.05), 145 of which were down-regulated (Fig. 3a and Supplementary Table 1), which indicates that Zbtb11 functions as a transcriptional activator. None of these genes were DE when comparing 4OHT-treated *Zbtb11^wt/wt^ R26^wt^* cells with EtOH-treated cells, indicating that their deregulation was not a result of the 4OHT treatment. To verify our RNA-seq results through a different method, we performed qRT-PCR for a panel of 20 genes spanning the entire range of effect sizes, and the results of the two methods were strongly correlated (Fig. 3b).

**Fig. 3.**
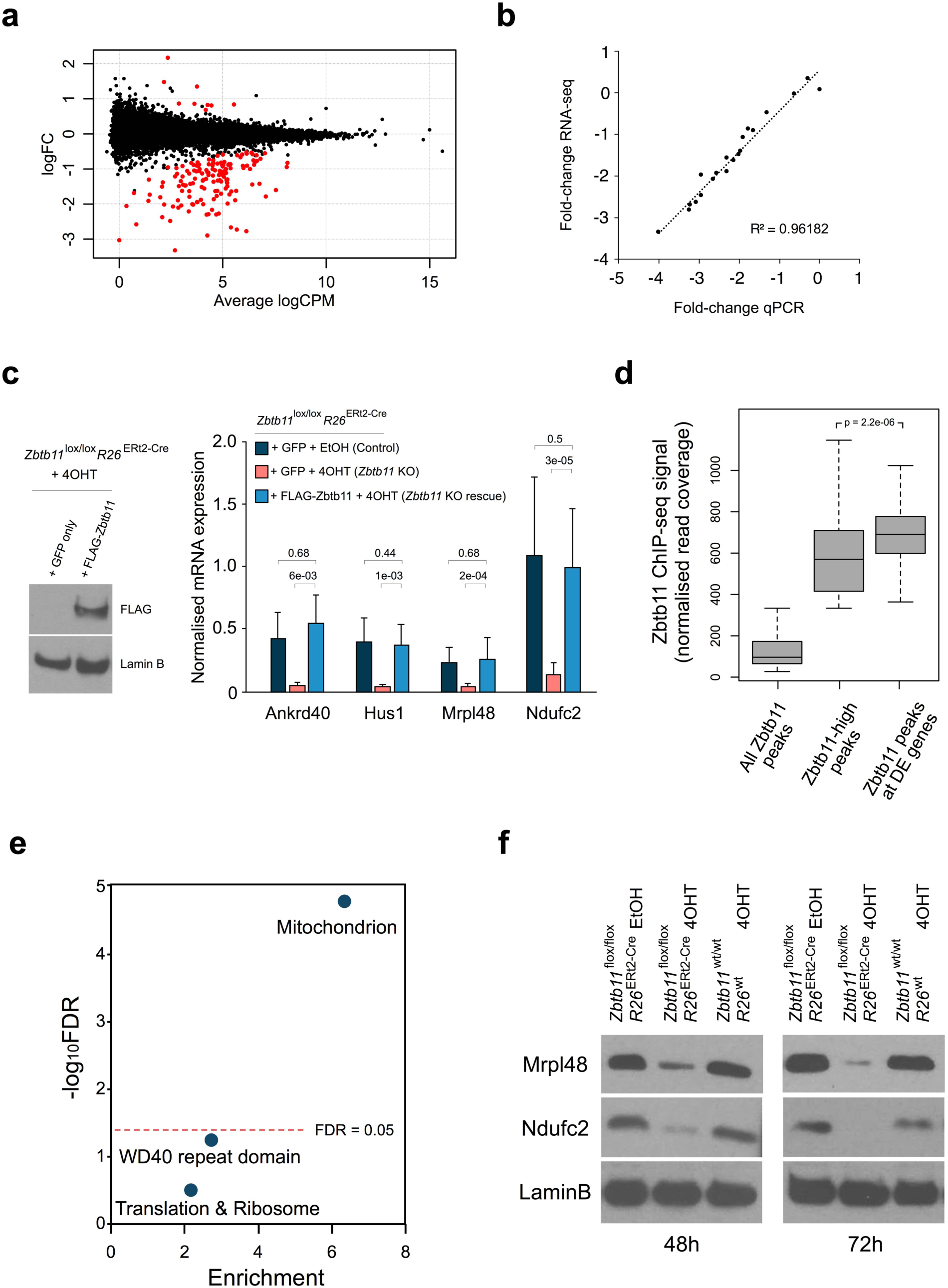
Zbtb11 supports the expression of a subset of nuclear encoded mitochondrial proteins. (a) Whole-transcriptome analysis by RNA-seq comparing *Zbtb11*^lox/lox^ *Rosa26*^ERt2-Cre^ cells treated with either EtOH (Control) or 4OHT (*Zbtb11* KO) for 48 hours. Plot shows log_2_-transformed fold-change (logFC) against transcript abundance (mean counts per million, logCPM) for all genes expressed in ESCs. Differentially expressed (DE) genes (FDR < 0.05) are shown in red. (b) Fold-change values obtained by RNA-seq for 20 genes plotted against fold-change values obtained by qRT-PCR for the same genes (mean of 3-5 biological replicates). (c) Zbtb11 expression rescue experiment. *Zbtb11*^lox/lox^ *Rosa26*^ERt2-Cre^ cells were transfected with a plasmid expressing either FLAG-Zbtb11–IRES–GFP, or GFP alone as control, and treated with either EtOH (Control) or 4OHT (*Zbtb11* KO). GFP-positive cells were FACS-sorted 48 hours post-treatment. Left panel - immunoblot of whole-cell lysates showing the expression of Zbtb11 protein was specifically restored. Right panel - qRT-PCR measuring the expression of a panel of genes down-regulated in *Zbtb11* KO cells. (d) Boxplot showing the distribution of Zbtb11 ChIP-seq signal strength among peaks found at promoters of DE genes, compared to Zbtb11-high peaks, and the entire peak set. Outlier values in the distribution of the entire Zbtb11 peak set are not shown as these make up the Zbtb11-high group (see also Supplementary Fig.1f). P-value (Wilcoxon rank sum test) is shown above. Note that peaks found at promoters of DE genes are overall significantly stronger than the rest. (e) DAVID functional annotation clustering analysis carried out on the genes differentially expressed in *Zbtb11* KO indicates one significant (FDR < 0.05) cluster of annotation terms, which is related to mitochondria. Significance (-log_10_FDR) is plotted against enrichment of the annotation terms associated with the genes. (f) Immunoblots for two mitochondrial proteins (Mrpl48 and Ndufc2) for which transcripts were found to be down-regulated in *Zbtb11* KO cells. LaminB was used as loading control.

To verify that the transcriptional changes are specific to Zbtb11 depletion, we performed experiments in which we rescued the expression of Zbtb11 in KO cells by ectopically expressing its cDNA. We transfected *Zbtb11^lox/lox^ R26^ERt2-Cre^* cells with a plasmid expressing either FLAG-Zbtb11– IRES–GFP, or GFP alone as control, and treated them for 48 hours with either EtOH or 4OHT. We subsequently FACS-sorted GFP-positive cells to isolate successfully transfected cells, confirmed specific expression of the FLAG-Zbtb11 protein by immunoblotting (Fig. 3c), and subsequently performed qRT-PCR to measure the expression of a panel of genes found to be down-regulated in the RNA-seq dataset. As seen in Fig. 3c, the expression of these genes was specifically rescued in the 4OHT-treated cells transfected with FLAG-Zbtb11 but not in 4OHT-treated cells transfected with GFP only. These results confirm the specificity of our inducible *Zbtb11* KO ESC line and show that the transcriptional changes detected by RNA-seq are indeed the result of Zbtb11 depletion.

Integration of the RNA-seq and ChIP-seq datasets revealed that 147 of the 154 DE genes (95%) have Zbtb11 peaks at their promoters, and that these binding sites are overall the strongest in the data set (Fig. 3d). This firmly indicates that Zbtb11 is directly regulating these genes, and that strength of binding is a good indicator of whether a Zbtb11 binding event is functional or not.

To determine which processes are controlled by Zbtb11, we performed functional enrichment analyses using the list of DE genes. This identified only one significant cluster of annotation terms (FDR < 0.05), which is defined by mitochondria and had an enrichment score of 6.12 (Fig. 3e and Supplementary Table 2). To investigate more thoroughly the association between Zbtb11 and genes with mitochondrial functions, we cross-referenced the DE genes with the MitoMiner database (Smith & Robinson, 2016), and found that 27% encode proteins with mitochondrial function, which represents an enrichment of 4.5 relative to the genes tested. All DE genes were nuclear-encoded, and they did not include any of the factors previously implicated in the regulation mitochondrial biogenesis and function, such as NRF-1, NRF-2/GABP, or members of the PGC-1, PPAR, or ERR families.

Finally, to verify that the changes observed at the mRNA level are reflected at the protein level, we carried out immunoblots on whole cell lysates from *Zbtb11^lox/lox^ R26^ERt2-Cre^* and *Zbtb11^wt/wt^ R26^wt^* cells treated with either EtOH or 4OHT for 48 and 72 hours. As seen in Fig. 3f, the mitochondrial proteins Mrpl48 and Ndufc2, encoded by DE genes, were already strongly down-regulated 48 hours post-treatment. These changes were not detected in 4OHT-treated *Zbtb11^wt/wt^ R26^wt^* cells, indicating they are not caused by 4OHT treatment alone. Altogether these results show that Zbtb11 is directly regulating the expression of a subset of nuclear-encoded mitochondrial proteins.

### Zbtb11 controls the locus-specific recruitment of NRF-2/GABP to its target promoters

To understand how Zbtb11 controls its target genes, we sought to identify other factors that Zbtb11 might functionally interact with. To this end we performed *de novo* motif discovery using the Zbtb11 ChIP-seq peakset, and separately the promoter regions of the 154 Zbtb11-dependent genes. Both approaches identified as the top hit the same motif, which closely matched that recognised by the ETS-domain protein GABPa (Fig. 4a), and which was firmly centred around the transcription start sites (TSS) (Fig. 4b). In addition, the consensus motif for Nuclear Respiratory Factor 1 (NRF-1) binding was also found to be enriched at the promoters of DE genes, but to a lesser extent and without a clear preference for the TSS (Fig. 4b). GABPa is the DNA binding subunit of the multimeric transcription factor GABP (GA-Binding Protein) also known as Nuclear Respiratory Factor 2 (NRF-2), which alongside NRF-1 plays essential roles in regulating mitochondrial biogenesis and function. Our motif analyses therefore suggested that Zbtb11 may regulate its target genes and mitochondrial activity through functional interactions with NRF-1 and/or NRF-2.

**Fig. 4.**
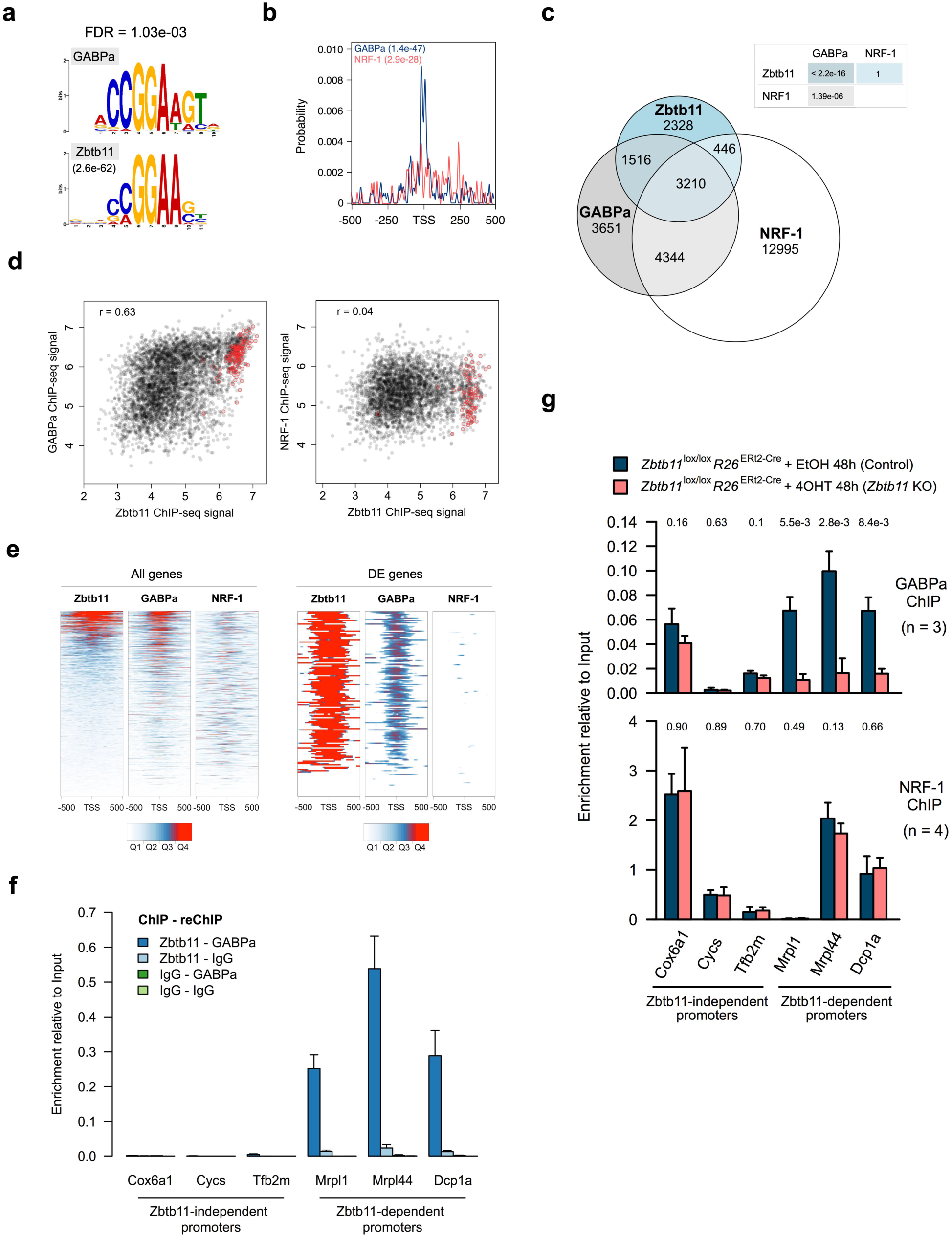
Zbtb11 controls the locus-specific recruitment of NRF-2. (a) Sequence logos for de novo Zbtb11 motif identified using (MEME ChIP E-value indicated) aligned to previously determined GABPa motif. FDR for the motif match in the database search is indicated above. (b) Probability distribution for GABPa and NRF-1 motifs in the promoter regions of Zbtb11-dependent genes. Enrichment p-values are given for each motif. (c) Genome-wide overlap between ChIP-seq peaks identified in mouse ES cell for Zbtb11, NRF-1 and GABPa. p-values for pairwise hypergeometric tests for overlap significance (see Methods) are shown in the table. (d) Plots of ChIP-seq signal (normalised read coverage) at overlapping peaks (black dots). peaks found at promoters of Zbtb11-dependent genes are circled in red. (e) Zbtb11, NRF-1 and GABPa ChIP-seq signal (quantiles of normalised read coverage) at the promoters of all genes expressed in mouse ESCs (left panels, n = 14,362), and at the promoters of Zbtb11-dependent genes (right panels, n = 154). (f) Sequential ChIP enrichment values at a panel of Zbtb11-dependent and -independent promoters in wild type mouse ESCs, using the indicated combinations of ChIP and reChIP andtibodies (mean and SD of three replicates). (g) GABPa and NRF-1 ChIP enrichment values at a panel of Zbtb11-dependent and -independent promoters, in control and Zbtb11 KO cells. P-values for two-tailed t-tests comparing control to Zbtb11 KO samples are shown above each binding site.

To test whether these motif analyses were corroborated by experimental evidence, we integrated our Zbtb11 ChIP-seq data with previously published ChIP-seq datasets for GABPa (Hartl *et al*, 2019) and NRF-1 (Domcke *et al*, 2015) in mouse ESCs. This revealed significant genome-wide overlap between Zbtb11 and GABPa peaks (p-value < 2.2e-16), but not between Zbtb11 and NRF-1 peaks (Fig. 4c). 63% of Zbtb11 peaks overlap with GABPa, but this fraction increases to 97% among Zbtb11-high peaks. While Zbtb11 overlaps with NRF-1 mainly at sites that are also occupied by GABPa, there is extensive NRF-1-independent overlap between Zbtb11 and GABPa. In agreement with previous studies showing that NRF-1 and NRF-2 share some common target genes, we find significant genome-wide overlap between NRF-1 and GABPa peaks, including at many Zbtb11-independent sites (Fig. 4c). Illustrative of the different locus-specific associations between Zbtb11, NRF-2 and NRF-1, these factors co-occupy the promoters of some mitoribosome subunit genes, but Zbtb11 does not bind classical NRF-1 and NRF-2 targets such as the promoters of cytochrome c, mitochondrial transcription factors TFB or cytochrome c oxidase subunit genes (Supplementary Fig. 3). At the promoters of DE genes, virtually all (98%) Zbtb11 peaks are also bound by GABPa, but only half of them are bound by NRF-1.

Analysing the strength of binding at shared sites, we found strong positive correlation genome-wide between Zbtb11 and GABPa (r = 0.63, p-value = 2.516e-16), with the peaks at DE genes overwhelmingly showing strong affinity for both Zbtb11 and GABPa (Fig. 4d, left panel). By contrast, we found no correlation between Zbtb11 and NRF-1 ChIP-seq signal (r = 0.04, p-value = 0.012) either genome-wide or at DE genes (Fig. 4d, right panel).

A promoter-centric analysis focusing on the ChIP-seq signal across the 1kb region around the TSS of all genes expressed in mouse ESCs (n = 14,362), revealed a positive correlation between Zbtb11 and GABPa (r = 0.49) and between GABPa and NRF-1 (r = 0.34), but not between Zbtb11 and NRF-1 (r = 0.10) (Fig. 4e, left panel). When considering only the promoters of the DE genes, Zbtb11 is very strongly correlated with GABPa (r = 0.74), but only weakly correlated with NRF-1 (r = 0.27) (Fig. 4e, right panel). Our analyses therefore strongly indicate that Zbtb11 functionally interacts with NRF-2, but do not support an interaction between Zbtb11 and NRF-1.

The strong positive correlation between Zbtb11 and GABPa binding suggested that these factors may bind cooperatively at their shared targets. Alternatively, Zbtb11 and GABPa could bind independently to the same genomic sites, which could take place on the same or on different alleles in the cell population, their positive correlation potentially being driven by their ability to recognise the same DNA binding motif. To distinguish between these possibilities, we performed sequential ChIP (ChIP-reChIP) with antibodies to Zbtb11 and GABPa. As seen in Fig. 4f, chromatin fragments bound by Zbtb11 are also strongly bound by GABPa, showing that Zbtb11 and GABPa bind simultaneously to their shared genomic sites. To explore the potential cooperative binding of these factors, we performed GABPa and NRF-1 ChIP in control and *Zbtb11* KO cells. This revealed that Zbtb11 depletion strongly impairs GABPa recruitment to shared Zbtb11/GABPa sites, but not to Zbtb11-independent sites (Fig. 4g). By comparison, NRF-1 recruitment was not affected by Zbtb11 depletion, either at shared Zbtb11/NRF-1 sites or at Zbtb11-independent sites (Fig. 4g). Zbtb11 therefore specifically controls the locus-specific recruitment of NRF-2 to its target promoters. Although Zbtb11 also shares some target genes with NRF-1, their association with these promoters appears to take place through independent mechanisms.

Altogether our results strongly indicated that Zbtb11 cooperates with NRF-2 to regulate mitochondrial function.

### Zbtb11 depletion results in impaired mitochondrial respiration

To test whether mitochondrial function is affected in *Zbtb11* KO cells, we first investigated the status of the mitochondrial membrane potential (MMP) by staining live cells with TMRE (tetramethylrhodamine, ethyl ester), a cationic fluorophore that permeates the plasma membrane and accumulates in the mitochondrial matrix in MMP-dependent manner. As can be seen in Fig. 5a (left panel) the TMRE staining decreased progressively after Zbtb11 depletion. This was not the result of fewer mitochondria, as the ratio of mitochondrial to nuclear DNA was the same in control and *Zbtb11* KO cells (Supplementary Fig. 4a). These results indicate that mitochondria gradually depolarise in *Zbtb11* KO cells, and show that Zbtb11 is required for the maintenance of functional mitochondria but not for mitochondrial replication.

**Fig. 5.**
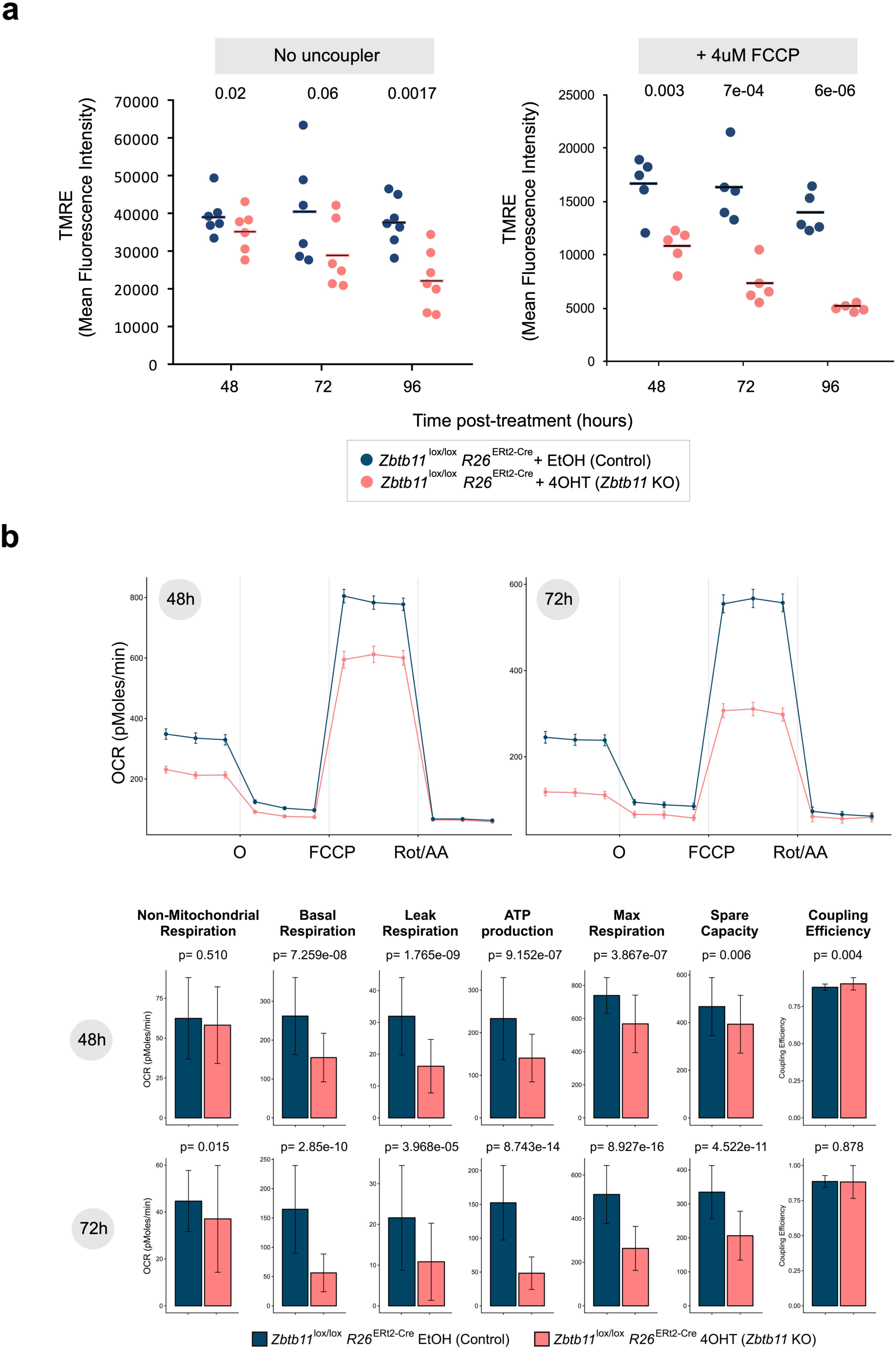
Zbtb11 depletion results in impaired mitochondrial respiration. (a) Changes in mitochondrial membrane potential (MMP) caused by Zbtb11 depletion. Control and *Zbtb11* KO cells were stained with TMRE (Tetramethylrhodamine, ethyl ester) at the indicated time following KO induction. Mean fluorescence intensity obtained by flow cytometry and their means are shown in samples without (left) and with 4uM FCCP (right). P-values for paired two-tailed t-tests are shown above. See also Supplementary Fig. 5b for full FCCP titration curves. (b) Decreased mitochondrial respiration in *Zbtb11* KO cells. *Zbtb11*^lox/lox^ *Rosa26*^ERt2-Cre^ cells were treated with either EtOH or 4OHT for 48 or 72 hours, and oxygen consumption rates (OCR) were measured using the Seahorse platform before and after the sequential addition of the ATP synthase inhibitor oligomycin (O), the mitochondrial uncoupler FCCP, and a mix of rotenone (complex I inhibitor) and antimycin A (complex III inhibitor) (Rot/AA). Upper panels - OCR values adjusted for gDNA content (measured by qPCR and used as a proxy for cell number), showing mean ± SEM of 8 (48h) and 7 (72h) biological replicates, and a total of 44-48 measurements (technical x biological). Lower panels – mitochondrial respiration parameters (mean ± SD) in control and *Zbtb11* KO cells, calculated from OCR measurements (see Methods). P-values (Wilcoxon rank sum test) are indicated above each pair.

To test the ability of *Zbtb11* KO cells to maintain their MMP in more stressful circumstances than the resting state, e.g. in situations when ATP demands suddenly increase, we challenged them with increasing concentrations of the mitochondrial uncoupler FCCP (Carbonyl cyanide 4-(trifluoromethoxy)phenylhydrazone). FCCP allows protons to flow back into the mitochondrial matrix, short-circuiting the MMP and stimulating the ETC complexes, which respond by increasing their activity to restore the cross-membrane proton gradient (Brand & Nicholls, 2011). We stained control and *Zbtb11* KO cells with TMRE in the presence of increasing concentrations of FCCP and subsequently analysed them by flow cytometry. As seen in Fig. 5a (right panel) and Supplementary Fig. 4b (left panel), the MMP in *Zbtb11* KO cells showed increased sensitivity to FCCP, which gradually intensified following Zbtb11 depletion. These effects were not observed in *Zbtb11^wt/wt^ R26^wt^* cells treated with 4OHT (Supplementary Fig. 4b, right panel).

The inability of *Zbtb11* KO cells to maintain polarised mitochondria could be caused either by decreased respiration or by an increase in proton leak or ATP demand. To determine whether respiration is impaired in Zbtb11-deficient cells, we used the Seahorse platform to measure oxygen consumption rates (OCR), which in combination with sequential addition of specific mitochondrial inhibitors (oligomycin, FCCP and a mixture of rotenone and antimycin A) allows the derivation of several mitochondrial respiration parameters, as previously described (Brand & Nicholls, 2011) (see Methods). This revealed that respiration is already significantly impaired 48 hours post-KO induction, and it further deteriorates 24 hours later (72 hours post-KO induction) (Fig. 5b). Basal respiration was the most affected, both its constituent parameters (leak respiration and ATP synthesis) showing the largest decrease of all parameters measured, which shows that mitochondrial depolarisation in *Zbtb11* KO cells is caused by deficiencies in the ETC and not by increased proton leak or ATP turnover. The coupling efficiency between respiration and ATP synthesis did not change in *Zbtb11* KO cells at any of the time points assessed, which indicates that Zbtb11 depletion has no direct effect on the proton leak, and that the reduction in leak respiration is just a consequence of a reduced proton flux through the system. The maximal respiratory capacity was also decreased in *Zbtb11* KO cells –although less affected than basal respiration to begin with, it subsequently deteriorated significantly, in agreement with the observed increasing sensitivity of the MMP to the uncoupling agent FCCP (see Fig. 5a, right panel).

Altogether these results show that *Zbtb11* KO cells have a deficient OXPHOS system, which fails to maintain the MMP and thus to supply an adequate proton-motive force, both at resting state and when forced to function at maximum capacity. Importantly, the decline of mitochondrial functions in our experimental system closely followed the down-regulation of Zbtb11-dependent mitochondrial proteins, but preceded any overt signs of cell death or cell cycle arrest.

### Zbtb11 controls complex I and mitoribosome biogenesis

To determine more specifically how Zbtb11 depletion affects mitochondrial respiration, we performed pathway mapping and over-representation analysis using the list of Zbtb11-dependent genes with known or predicted mitochondrial functions. This identified ‘complex I biogenesis’ and ‘mitochondrial translation’ as the most likely affected processes (FDR < 0.05, Fig. 6a and Table S3). There were four genes involved in the biogenesis of respiratory complex I, encoding one core subunit (Ndufs7), two accessory subunits (Ndufa12 and Ndufc2), and one assembly factor (Ndufaf1) (Fig. 6a and Table S3). There were eight DE genes involved in mitochondrial translation, all of which encode subunits of the mitochondrial ribosome (Fig. 6a and Table S3). All DE genes that mapped to mitochondrial pathways were down-regulated, and the effect size at *Ndufc2* and *Ndufaf1* was particularly high (5.5- and 4-fold, respectively, see Fig. 6a). This acute depletion was also reflected at the protein level, as Ndufc2 was virtually undetectable by immunoblotting 72 hours after Zbtb11 KO induction (see Fig. 3f). These results suggested that *Zbtb11* KO cells may be defective in the synthesis and assembly of respiratory complex I. In addition, these cells may also have a compromised mitoribosome, which would cause a general defect in mitochondrial translation, and therefore have a knock-on effect on the synthesis of all OXPHOS complexes with mtDNA-encoded subunits (I, III, IV and V).

**Fig. 6.**
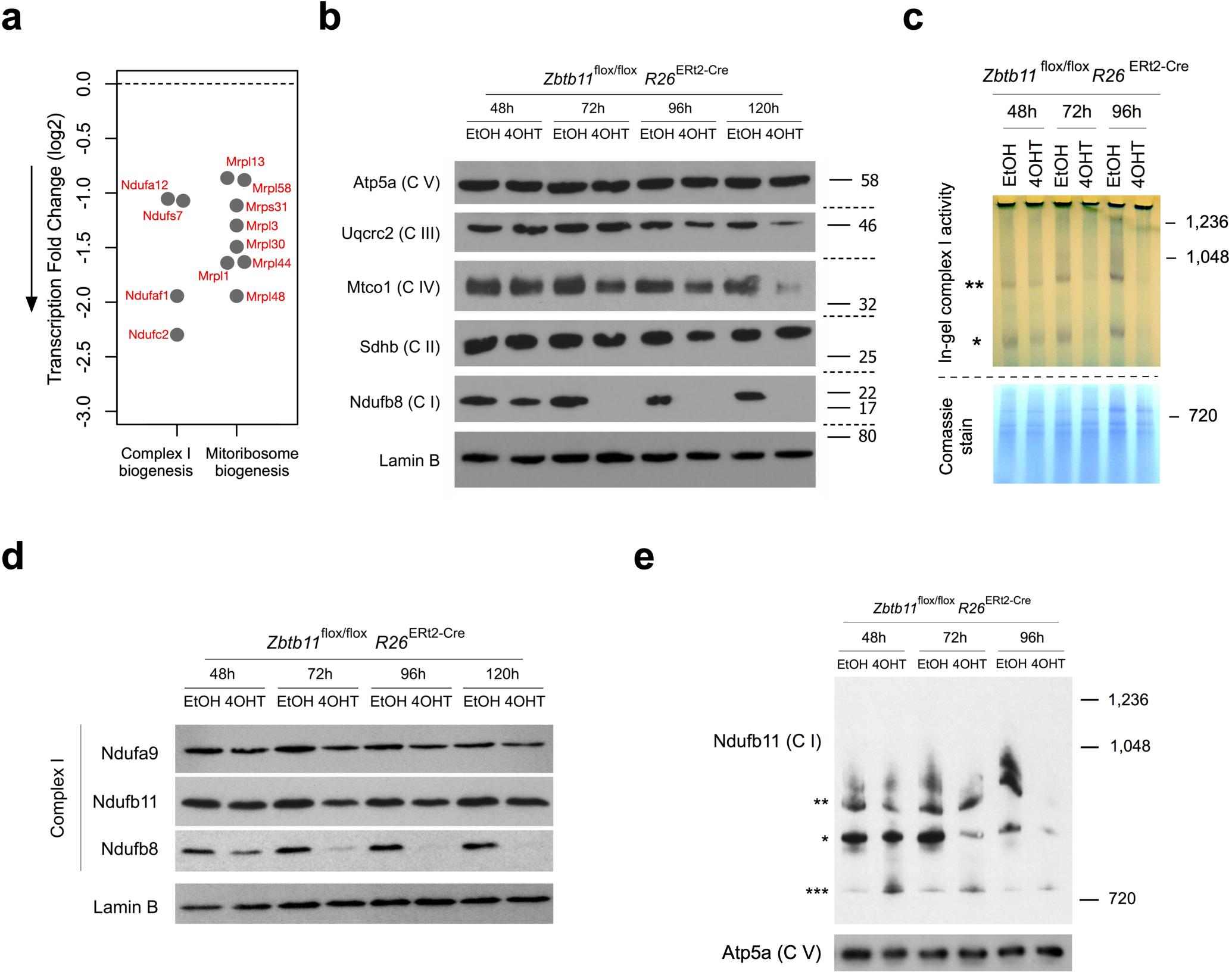
Zbtb11 controls assembly and stability of respiratory complex I. (a) Differentially expressed genes mapping to enriched mitochondrial pathways. (b) Immunoblots of whole cell lysates from control and *Zbtb11* KO cells, probing stability markers of all five OXPHOS complexes. (c) Blue native polyacrylamide gel electrophoresis (BN-PAGE) of mitochondrial extracts from control and *Zbtb11* KO cells. Left panel – in-gel complex I activity assay. Purple bands (* complex I; ** complex I-containing supercomplexes) develop following the incubation of the gel in a substrate solution (see Methods), their intensity providing a measure of complex I activity present in the mitochondrial extract. Note the reduction of complex I activity 48 hours post-KO induction and its near-absence from 72 hours onwards. Right panel – following the in-gel activity assay, the gel was stained with Comassie to reveal the protein standards and verify equal loading. (d) SDS-PAGE and immunobloting of complex I subunits in whole cell lysates of control and *Zbtb11* KO cells showing that not all complex I subunits are down-regulated. Lamin B is shown as loading control. (e) Immunoblot of native mitochondrial extracts separated by BN-PAGE. Note the reduction in complex I holocomplexes (*) and supercomplexes (**) and the increase in a smaller Ndufb11-containing assembly intermediate (***). Complex V detected with anti-Atp5a antibodies is shown as loading reference.

To assess the integrity of respiratory complexes in *Zbtb11* KO cells we carried out immunoblotting with antibodies against specific subunits that become degraded when their respective OXPHOS complex is unstable or fails to assemble (Lake *et al*, 2017; Maio *et al*, 2016, 2014). We found a striking down-regulation of complex I integrity marker Ndufb8 (Fig. 6b), despite the fact its transcript is expressed at normal level (fold-change 1.006, adjusted p-value = 0.99). The integrity markers for complex III (Uqcrc2) and complex IV (Mtco1) were also down-regulated, but only after prolonged Zbtb11 depletion (Fig. 6b). By comparison, the expression of complex II and V markers remained unchanged at all times (Fig. 6b). These results indicated complex I biogenesis is specifically affected in the immediate aftermath of Zbtb11 depletion, in agreement with the prediction by the differential gene expression analysis.

Blue native polyacrylamide gel electrophoresis (BN-PAGE) of mitochondrial extracts, followed by in-gel activity assays with a NADH substrate, showed that complex I activity is indeed reduced 48 hours post-*Zbtb11* KO induction, and becomes undetectable from 72 hours onwards (Fig. 6c). Further immunoblotting experiments showed that not all complex I subunits are degraded (Fig. 6d), indicative of an assembly defect rather than degradation of the entire holocomplex. To assess this, we separated mitochondrial extracts by BN-PAGE and subsequently performed immunoblotting with antibodies against Ndufb11, a subunit that is not degraded as a result of Zbtb11 depletion (Fig. 6d). This showed that 48 hours post-*Zbtb11* KO induction the holocomplex was already reduced, while a smaller Ndufb11-containing subcomplex was increased, consistent with an assembly defect (Fig. 6e). 72 hours post-KO induction, mitochondria were almost completely depleted of complex I holocomplex, and at 96 hours both the holocomplex and the subcomplex were barely detectable with Ndufb11 antibodies (Fig. 6e), indicating the subcomplex also becomes unstable or fails to assemble. Altogether, these data show that Zbtb11 depletion leads to a severe complex I assembly defect and complete loss of complex I activity, strongly indicating that the reduction in mitochondrial respiration that follows Zbtb11 depletion (see Fig. 5b) is underpinned to a significant extent by complex I insufficiency.

To understand whether the impaired biogenesis of respiratory complexes in *Zbtb11* KO cells is underpinned by a mitoribosome defect, we first tested if pharmacological inhibition of the mitoribosome mimics the respiratory complexes defect that follows Zbtb11 depletion. Blocking mitochondrial translation with chloramphenicol in otherwise untreated *Zbtb11^lox/lox^ R26^ERt2-Cre^* cells resulted in complete depletion of complexes I, III and IV within 48 hours of treatment, while complexes II and V remained unaffected (Fig. 7a). This shows that mitochondria-encoded subunits of complexes I, III and IV are rapidly turned over, leading to the collapse of these complexes 48 hours after their synthesis is blocked. This phenotype is largely recapitulated in 120 hours *Zbtb11* KO (Fig. 6b and Supplementary Fig. 5), indicating that mitochondrial translation may become impaired 72 hours post-*Zbtb11* KO induction. To test this possibility, we compared the dynamics of mitoribosome-dependent synthesis of respiratory complexes in control and *Zbtb11* KO cells. We first depleted complexes I, III and IV by treating cells with chloramphenicol for 48 hours, after which we removed the mitochondrial translation block, allowing it to resume, and subsequently monitored the regeneration of the respiratory complexes (Fig. 7b). By synchronising the chloramphenicol and EtOH/4OHT treatments in *Zbtb11^lox/lox^ R26^ERt2-Cre^* cells, we allowed mitochondrial translation to resume either 48 or 72 hours post-*Zbtb11* KO induction (Fig. 7b). Upon release from the mitochondrial translation block, control cells resumed synthesis of mitochondria-encoded proteins such as Mtco1, and regenerated all respiratory complexes within 48 hours (Fig. 7c). By contrast, *Zbtb11* KO cells were impaired in their ability to synthesise complexes I, III and IV (Fig. 7c). While synthesis of complex III and IV was impaired but not fully arrested, complex I biogenesis was completely blocked and never recovered after chloramphenicol removal. These results are consistent with a partial impairment of mitochondrial translation leading to reduced complex III and IV synthesis, but which in the case of complex I is compounded or fully masked by the specific and more severe biogenesis defect caused by the down-regulation of complex I subunits directly controlled by Zbtb11 (see Supplementary Table 3).

**Fig. 7.**
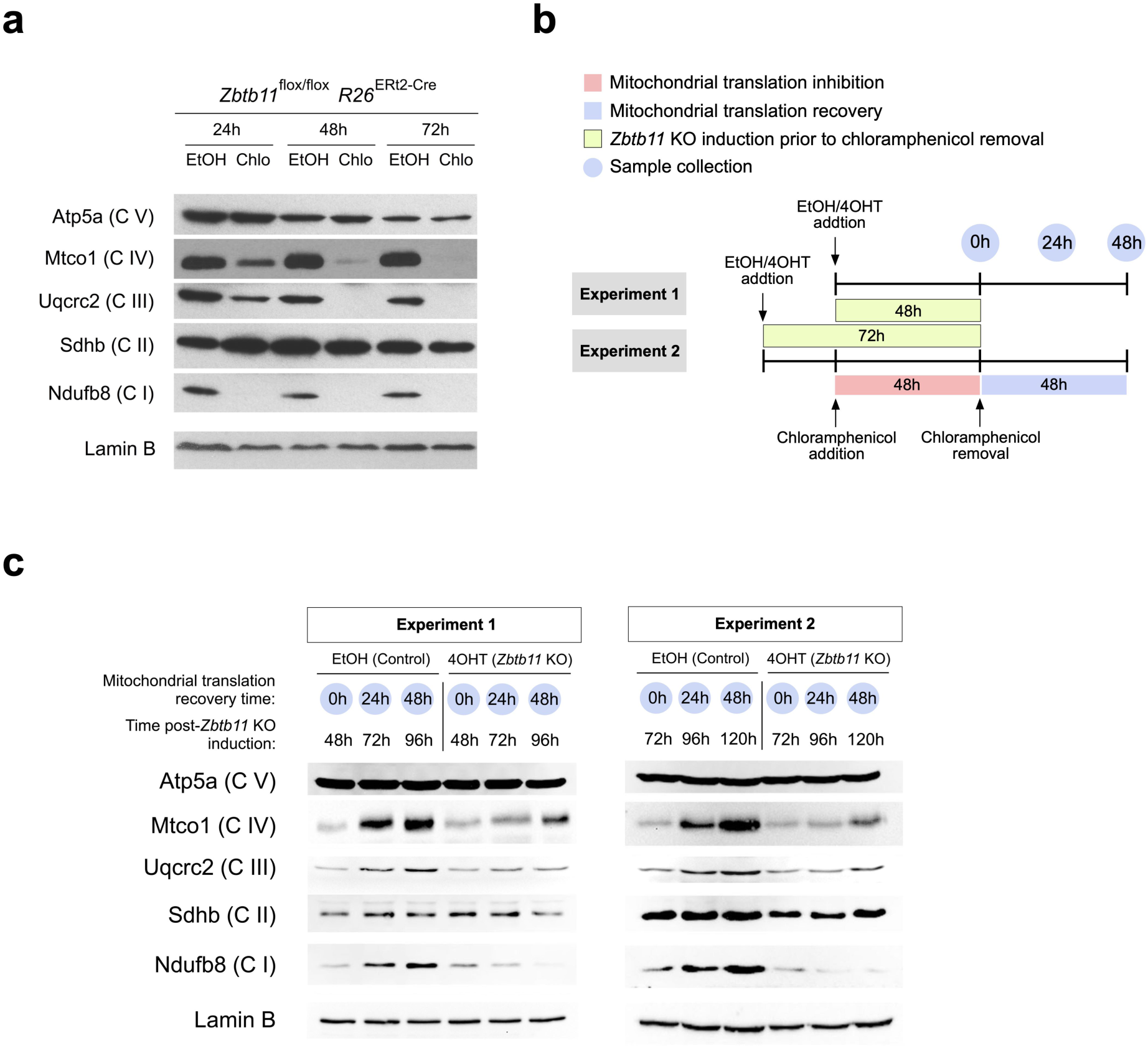
Zbtb11 is required for the biogenesis of respiratory complexes I, III, and IV. (a) SDS-PAGE and immunoblotting of whole cell extracts following inhibition of mitochondrial translation with chloramphenicol. Samples from cells treated with ethanol (carrier) were used as reference. Note that complexes I, III and IV are sensitive to the mitochondrial translation block. Lamin B is shown as loading control. (b) Diagram of the experimental design used to compare *de novo* synthesis of respiratory complexes in control and *Zbtb11* KO cells. Fully assembled complexes I, III and IV were depleted from *Zbtb11*^lox/lox^ *Rosa26*^ERt2-Cre^ cells by inhibiting mitochondrial translation for 48 hours (as in Fig. 7a). Mitochondrial translation was then allowed to resume by removing the chloramphenicol treatment, and samples for immunoblotting were collected at 24 hour intervals in order to monitor the regeneration of respiratory complexes. The chloramphenicol treatment was synchronised with the addition of 4OHT (or EtOH as control), so that translation was allowed to resume either 48 hours (experiment version 1) or 72 hours (experiment version 2) post-*Zbtb11* KO induction, thus allowing to compare *de novo* synthesis of respiratory complexes in control and *Zbtb11* KO cells. (c) Immunoblotting analyses of representative experiments comparing *de novo* synthesis of respiratory complexes I, III and IV, in control and *Zbtb11* KO cells, as described in Fig. 7b. Note that 48 hour *Zbtb11* KO cells can synthesise complexes III and IV but not complex I, while 72 hour Zbtb11 KO cells are not able to synthesise any of the three complexes sensitive to mitochondrial translation inhibition.

In conclusion, Zbtb11 specifically controls complex I biogenesis by promoting the expression of key accessory and assembly subunits, while also exerting – albeit to a lesser extent – broader control over the synthesis of respiratory complexes, most likely through its role in mitoribosome biogenesis.

### Mutations associated with intellectual disability destabilise the Zbtb11 protein and impair complex I biogenesis

In order to investigate the relevance of our findings to the aetiology of *ZBTB11*-associated intellectual disability, we modelled the mutations associated with this disease in our experimental system. Two *ZBTB11* missense mutations have been identified in families with hereditary intellectual disability, both of which lead to single amino acid substitutions of key histidine residues (H729Y and H880Q) (Fattahi *et al*, 2018, 11) in C2H2 zinc finger motifs that are conserved between mouse and humans. We induced *Zbtb11* KO and rescued its expression by ectopically expressing either wild type or mutant FLAG-Zbtb11 cDNA (H729Y or H880Q). The cDNA was linked to an IRES-GFP reporter, which allowed us to use flow cytometry to sort successfully transfected cells with the same level of cDNA expression. As seen in Fig. 8a, although the wild type and the two mutant FLAG-Zbtb11 cDNA were expressed at comparable levels, only wild type Zbtb11 was able to fully rescue the expression of complex I and mitoribosome biogenesis genes. Complex I biogenesis genes *Ndufc2* and *Ndufaf1* were particularly affected by the Zbtb11 mutations, their transcripts being downregulated 2-fold when Zbtb11 expression was rescued with H880Q and H729Y mutants, respectively. By comparison, the maximum effect size among the mitoribosome genes was 1.6-fold down-regulation as a result of mutating Zbtb11.

**Fig. 8.**
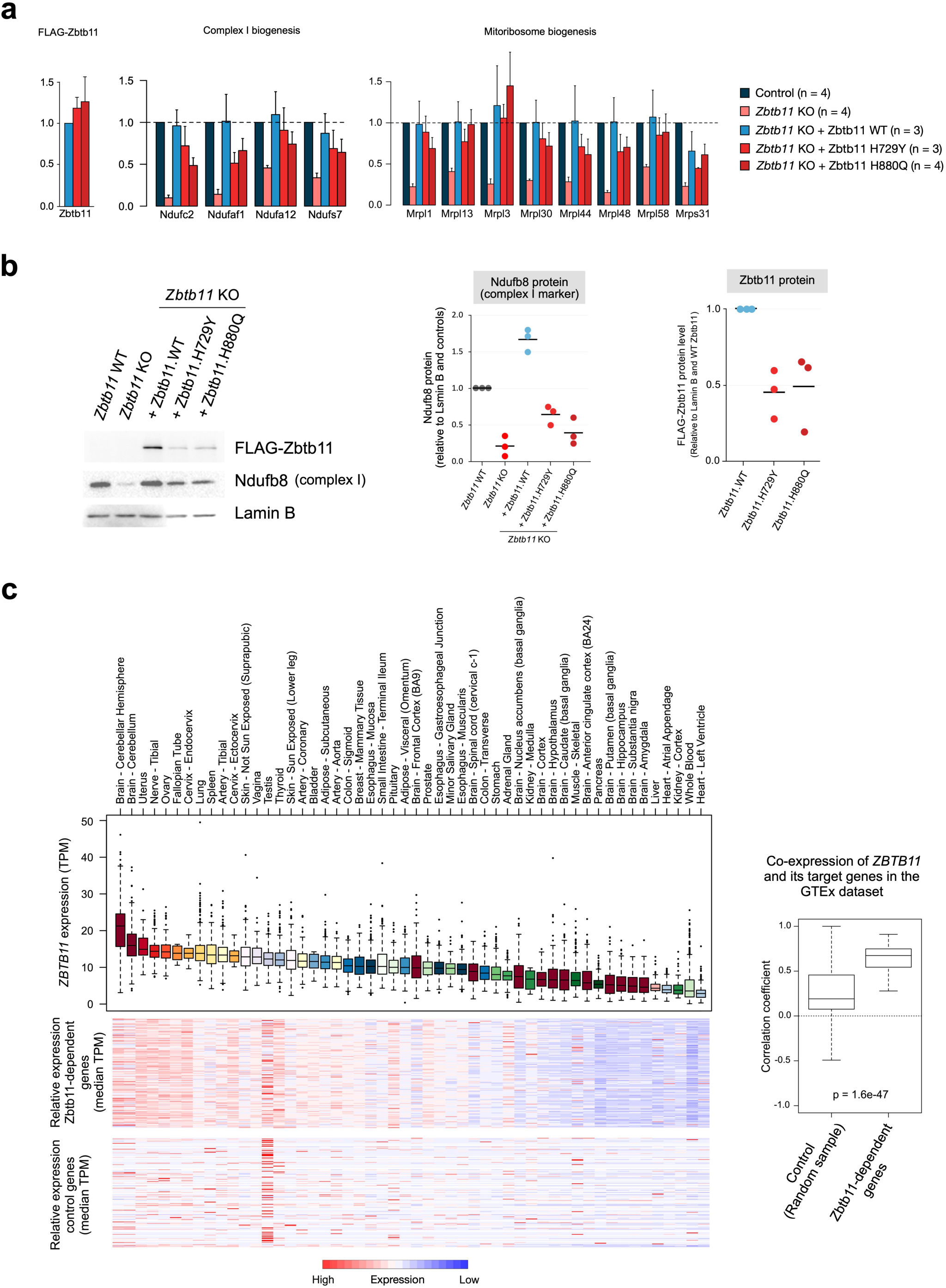
Mutations associated with intellectual disability destabilise the Zbtb11 protein and impair complex I biogenesis. (a) qPCR quantification of transcripts in control and *Zbtb11* KO cells rescued with either wild type (WT) or mutant Zbtb11 cDNA. *Zbtb11*^lox/lox^ *Rosa26*^ERt2-Cre^ cells were treated with either EtOH (Control) or 4OHT (*Zbtb11* KO), and concomitantly transfected with a plasmid expressing either WT or the indicated mutant FLAG-Zbtb11 cDNA fused to an IRES-GFP reporter. 48 hours later successfully transfected cells were sorted based on GFP expression, gating on cells with intermediate fluorescence, thus selecting cells expected to have similar levels of cDNA expression. (b) Immunoblot of whole cell lysates of *Zbtb11* KO cells rescued with either WT or mutant Zbtb11 cDNA. Cells treated and sorted as in (a), except collected 72 hours post-*Zbtb11* KO induction, were analysed by immunoblotting for the complex I stability marker (Ndufb8) and FLAG-Zbtb11 expression. Lamin B was used as loading reference. Left panel shows a representative image of the results, while the two panels on the right show the quantification of the blots from 3 biological replicates. (c) Boxplot to the left shows normalised expression of human *ZBTB11* across primary tissues sampled in the GTEx dataset. The heatmaps below represent the same tissues as in the boxplot and median expression values for the human orthologues of Zbtb11-dependent genes (upper panel), and for an equal number of randomly selected genes as control (lower panel). Boxplot to the right shows distribution of correlation coefficients for expression of *ZBTB11* and that of individual target genes across the GTEx tissues.

Immunoblotting with antibodies to the complex I stability marker Ndufb8, showed that the inability of mutant Zbtb11 to fully support transcription of complex I biogenesis genes results in reduced amounts of complex I (Fig. 8b). Surprisingly, immunoblotting with anti-FLAG antibodies revealed that mutant FLAG-Zbtb11 protein levels were also reduced approximately two-fold for both mutants (Fig. 8b), despite their respective cDNA transcripts being at least as abundant as wild type FLAG-Zbtb11 (Fig. 8a). This shows that the pathogenic mutations in the zinc finger motifs of Zbtb11 cause the protein to be unstable or mark it for degradation.

To frame our findings in the context of human tissue-specific gene expression, we integrated our results with the Genotype-Tissue Expression (GTEx) dataset (V8), which contains whole transcriptome data from 948 individuals across 52 different primary tissues. This revealed that while *ZBTB11* is ubiquitously expressed, it is most highly expressed in samples from the cerebellum (Fig. 8c) – in agreement with an important function for Zbtb11 in this tissue, also indicated by the fact that patients with *ZBTB11* mutations have cerebellar atrophy (Fattahi *et al*, 2018, 11). Of the 154 Zbtb11-dependent genes we identified in mouse ES cells, 149 were successfully matched to human orthologues. Their expression was highly correlated with *ZBTB11* (Fig. 8c), supporting the notion that the regulatory relationship of Zbtb11 with its target genes is conserved in mammals.

Altogether, our results indicate that the effect of human *ZBTB11* mutations is mediated at least in part by a reduction in the amount of Zbtb11 protein available to activate its target genes. This is likely to be particularly detrimental in tissues requiring higher Zbtb11 expression, such as the cerebellum, and may be further compounded by a defect in the ability of mutant Zbtb11 to recognise its cognate targets through the zinc finger motifs. Importantly, these mutations impair complex I biogenesis, indicating that the phenotype in affected patients may be, at least partly, the manifestation of a mitochondrial disease. This is supported by the phenotypical overlap with other mitochondrial diseases, which are also commonly characterised by cerebellar atrophy, ataxia and hypotonia (Lax *et al*, 2012).

## Discussion

There are currently approximately 700 genes known to be involved in the genetics of intellectual disability (ID) (Vissers *et al*, 2016), but it is estimated the total number may be greater than 2,500 (Musante & Ropers, 2014). Large-scale studies of consanguineous families with inherited ID have identified new candidate loci, including several genes encoding proteins with unknown function (Najmabadi *et al*, 2011; Harripaul *et al*, 2017). Determining the molecular and cellular functions of these proteins is a crucial step towards understanding the aetiology of the different forms of ID. Zbtb11 is a ubiquitously expressed transcription factor recently found to be mutated in two families with inherited ID (Harripaul *et al*, 2017; Fattahi *et al*, 2018), but which had remained largely uncharacterised. In this study we performed a systematic analysis of its cellular functions.

We found Zbtb11 is an essential factor required for cell proliferation and survival, which explains why only missense mutations, but no homozygous loss of function variants, have been identified so far in ID patients or in the Genome Aggregation Database (gnomAD) (Karczewski *et al*, 2019). Using an inducible KO system allowed us to perform timed deletion experiments and detect the early transcriptional changes at genes under direct Zbtb11 control. Zbtb11 appears to function as a transcriptional activator, contrasting with the majority of previously characterised ZBTB proteins, most of which have been reported to function as transcriptional repressors (Lee & Maeda, 2012).

A recent study reported that in zebrafish, Zbtb11 directly regulates p53 by binding to its gene promoter and repressing transcription (Keightley *et al*, 2017). Inspection of our ChIP-seq datasets also revealed binding of Zbtb11 at the promoter of p53 gene (*Trp53*), but *Trp53* was not deregulated in our RNA-seq experiment (fold-change = 0.93, adjusted p-value = 0.99) (Supplementary Fig. 6a). qRT-PCR analyses confirmed that *Trp53* transcripts do not change in *Zbtb11* KO cells, even after prolonged Zbtb11 depletion 72 hours post-KO induction (Supplementary Fig. 6b), leading us to conclude that Zbtb11 does not directly regulate the expression of p53 in mouse ESCs. The discrepancy between our results and the previous study may be due to species-specific differences. Alternatively, the zebrafish *Zbtb11* missense mutation found to upregulate p53 may act as a gain of function mutation that potentiates the transcriptional activating function of Zbtb11.

We find instead that in mammalian cells Zbtb11 promotes the expression of a select group of house-keeping genes that are significantly enriched in genes encoding mitochondrial proteins. Promoting the expression of these mitochondrial proteins has functional importance, as Zbtb11 depletion significantly impairs respiration and leads to mitochondrial depolarisation. These changes precede detectable increases in cell death, indicating they are not a consequence of apoptosis. Maintenance of mitochondrial function is therefore a key role of Zbtb11.

Mechanistically, we show that Zbtb11 is an integral part of the network of nuclear transcription factors which controls mitochondrial activity and biogenesis, being responsible for the recruitment of NRF-2/GABP to a subset of its target sites. Zbtb11 shares a large proportion of its genomic binding sites with NRF-2/GABP and our experiments established these two factors bind simultaneously at their common targets, where NRF-2/GABP recruitment is heavily dependent on Zbtb11. Importantly, this functional association is a signature of virtually all Zbtb11-high sites and promoters of Zbtb11-dependent genes, indicating that recruitment of NRF-2/GABP is part of the mechanism through which Zbtb11 controls its targets genes. NRF-2/GABP is a potent promoter activator, as recently illustrated by the finding that in a large proportion of glioblastomas the telomerase gene is reactivated by promoter mutations that generate novel NRF-2/GABP binding sites (Bell *et al*, 2015). The transcriptional activating role of Zbtb11 can therefore be explained by its ability to facilitate the recruitment of NRF-2/GABP to promoters.

Interestingly, despite their extensive genome-wide association, NRF-2/GABP also binds to a number of promoters, including some of its known classical targets, where its binding is not Zbtb11-dependent. This shows that NRF-2/GABP does not bind its targets through a uniform mechanism, but its recruitment is instead regulated in locus-specific manner. This may reflect the existence of locus-specific complexes in which NRF-2/GABP associates with different factors. Alternatively, Zbtb11 may facilitate promoter activation by regulating the local chromatin structure to increase TSS accessibility for NRF-2/GABP.

Zbtb11 was found to co-localise with NRF-1 at the promoters of several genes, the majority of which, however, are also bound by NRF-2/GABP. At these sites Zbtb11 specifically controls the recruitment of NRF-2/GABP but not NRF-1, showing that although NRFs overlap extensively genome-wide, their activity can be distinctly regulated even at shared promoters.

Zbtb11 controls the biogenesis of complex I by promoting the expression of nuclear-encoded complex I subunits with core (Ndufs7), accessory (Ndufc2 and Ndufa12) and assembly (Ndufaf1) roles, Ndufc2 and Ndufaf1 being particularly sensitive to Zbtb11 depletion (see Supplementary Table 1). An abundance of evidence from studies of human disease (Dunning *et al*, 2007; Lebon *et al*, 2007; Ostergaard *et al*, 2011) and *in vitro* experimental approaches (Stroud *et al*, 2016) show that inactivating either one of these factors individually leads to severe defects in complex I assembly, so their combined down-regulation in *Zbtb11* KO cells is expected to lead to the collapse of complex I – an outcome that is reflected in our results. Zbtb11 also directly controls the expression of mitoribosomal protein components, and its depletion eventually caused a broad defect in the biogenesis of respiratory complexes that partly recapitulates the pharmacological inhibition of mitochondrial translation. This effect, however, was not as pronounced as the specific impairment of complex I biogenesis, which may be due to the fact that the majority of the mitoribosome genes were not down-regulated to the same extent as the complex I genes *Ndufc2* and *Ndufaf1*.

In conclusion, Zbtb11 maintains the homeostasis of the OXPHOS system primarily by activating transcription at genes of nuclear-encoded complex I subunits, as well as by indirectly facilitating the expression of mitochondria-encoded subunits at the translational level.

Experimental modelling of the missense Zbtb11 mutations associated with ID showed that these have a destabilising effect on the Zbtb11 protein. Both mutations lead to the substitution of the second histidine residue in C2H2 zinc finger motifs, which is predicted to disrupt the folding of the motifs, and protein misfolding may accelerate its degradation. The overall effect of the mutations is therefore equivalent to a Zbtb11 knock-down, which could be further compounded by the fact that the mutant protein may be impaired in its ability to bind and activate target promoters. The complex I genes *Ndufc2* and *Ndufaf1* were particularly sensitive to the reduced Zbtb11 dosage, and consequently, Zbtb11 mutations lead to impaired complex I biogenesis. Our findings provide a rationale for the aetiology of this form of ID, as neuronal development, survival and activity, are heavily reliant on mitochondria for ATP production and for regulation of calcium and redox signalling. Mitochondria play key roles in axonal specification and growth (Ruthel & Hollenbeck, 2003) as well as in synaptic plasticity (Li *et al*, 2004), and their importance in neuronal development and survival is underlined by the fact that many mitochondrial disorders are characterised by neurological abnormalities. Of note, cerebellar atrophy and ataxia are prominent features in patients with mtDNA pathogenic mutations (Lax *et al*, 2012), and are also evident in patients with *ZBTB11* mutations. Moreover, at least two genes that we find to be directly regulated by Zbtb11, *NDUFA12* and *NDUFAF1*, are known to be mutated in mitochondrial diseases (Leigh Syndrome and cardioencephalomyopathy, respectively) (Ostergaard *et al*, 2011; Stroud *et al*, 2016). It is therefore likely that at least some of the phenotypical features reported in patients with *ZBTB11* mutations are manifestations of a mitochondrial disease.

## Data Accession

All high-throughput sequencing data in this study have been deposited in the Gene Expression Omnibus repository (GEO: GSE125047).

## Materials and Methods

### Cells

Mouse ES cell line E14 (Doetschman et al. 1987) and its derivatives were maintained on plates pre-coated with 0.2% gelatin (G1393, Sigma). Media was Knockout DMEM (10829018, Gibco), supplemented with foetal bovine serum (10%) (10270106, Thermo Fisher Scientific), non-essential amino acids (11140035, Gibco), glutamine (2mM), beta-mercaptoethanol, penicillin and streptomycin, and LIF (ESG1106, Merck-Millipore).

### CRISPR/Cas9-mediated genome editing

Guide RNA (gRNA) sequences were synthesised as single stranded oligos (Sigma-Aldrich), annealed and cloned in the vectors PX458 (pSpCas9(BB)-2A-GFP, Addgene plasmid # 48138) or PX461(pSpCas9n(BB)-2A-GFP, Addgene plasmid # 48140), both a gift from Feng Zhang (Ran et al. 2013). Recombinant plasmids were verified by Sanger sequencing and electroporated into E14 cells using the Nucleofector II platform (Lonza) (program A-013). 48 hours post-electroporation GFP-expressing cells were FACS-sorted and cloned by serial dilution in 96-well plates. Single cell colonies were allowed to grow for a week and subsequently screened by PCR. Colonies detected as recombinant were further analysed by Sanger sequencing, by PCR amplifying the targeted locus, cloning the PCR amplicons in the pCR2.1 TOPO plasmid (Thermo Fisher Scientific) and transforming 10-beta competent *E. coli* bacteria (New England Biolabs). Plasmid inserts from 20 bacterial colonies were sequenced for each ES cell colony to verify homozygosity.

### Generation of the *Zbtb11*^FLAG/FLAG^ E14 line

The *Zbtb11* ATG start codon was targeted using two gRNAs (GCCGGCGGCTATGTCAAGCG and CGCCTGTCAGTGGTAAGGAG) cloned in PX461, which co-expresses the Cas9 nickase and GFP. A donor template for homology-directed repair (HDR), that contained the modified ATG start site with the 3xFLAG tag flanked by two 800bp homologous arms, was also electroporated at the same time. Recombinant colonies were screened by PCR using the primers TGGGAGAAAGATGCTCTCCAT and CACCGTCATGGTCTTTGTAGTC. Primers TGGGAGAAAGATGCTCTCCAT and CCTCGCTTGACATAGCCGCC detect the wild type unmodified locus but not the *Zbtb11*^FLAG^ allele.

### Generation of the inducible *Zbtb11* KO line

In the first step introns 2 and 3 were targeted using the gRNAs TAGGATTAAGGAAAACATTG and GCTTCTAATACTCTGTGCAA, respectively, cloned in PX458 which co-expresses the wild type Cas9 nuclease and GFP. The plasmids encoding the two gRNAs were co-electroporated with a HDR template containing a 2.8kb region spanning *Zbtb11* exon 3 and flanking intronic sequences modified to include the loxP sequence alongside ectopic restriction sites at the regions targeted by the gRNAs (see also Fig. 2a). Successfully targeted colonies were detected by PCR (GATAGGGAGCCCTGCTCTCA and CCACAGCCTCCAAGTCTTCC) followed by restriction digest with EcoRI and BamHI (see also Supplementary Fig.2a). Following the isolation of a homozygous *Zbtb11*^lox/lox^ line, the ERt2-Cre transgene was inserted into the *Rosa26* locus using the gRNA ACTCCAGTCTTTCTAGAAGA cloned into PX458, and the pMB80 plasmid (a gift from Tyler Jacks, Addgene plasmid # 12168) (McLaughlin et al. 2007) as HDR template. Recombinant colonies were detected by nested PCR with the primers GGCCCAAATGTTGCTGGATA and AGAGCCTCGGCTAGGTAGGG in the first step, and AGGTTCTGCGGGAAACCATT and AGGTAGGGGATCGGGACTCT in the second step.

To induce deletion of *Zbtb11* exon 3, cells were treated with 4-hydroxytamoxyfen (T176, Sigma-Aldrich) by adding it to the media from a 1,000x stock dissolved in ethanol, to a final concentration of 250nM.

### Genomic and mitochondrial DNA qPCR

DNA was extracted using the DNeasy Blood & Tissue Kit (Qiagen). 20ng of total DNA was used in a qPCR reaction containing QuantiNova SYBR Green mix (Qiagen), which was ran on a StepOnePlus Real-Time platform (Thermo Fisher Scientific). To quantify gDNA the geometric mean of two primer pairs amplifying different genomic loci was used. The geometric mean of two separate mitochondrial loci was used to quantify the mtDNA. The number of functional *Zbtb11*^lox^ alleles remaining in the cell population after KO induction was quantified using an intron 3-specific primer and a primer that specifically recognises the intron 3 loxP site but not the intron 2 loxP site.

Cre-mediated recombination disrupts the binding site of the loxP-specific primer and therefore qPCR signal is no longer generated from the KO alleles. The signal for functional *Zbtb11*^lox^ alleles was normalised to the geometric mean of the two gDNA primer pairs.

### qRT-PCR

RNA was extracted using the RNeasy Mini kit (Qiagen). Exogenous ERCC RNA Spike-In Mix 1 (Thermo Fisher Scientific) was added at the cell lysis stage in a ratio of 1μl per 9 million cells. 1μg total RNA was used to synthesise cDNA with the ProtoScript II kit (New England Biolabs) using random primers in a 20μl reaction. The RT-PCR reaction was diluted to 100μl with ultrapure water after which 1μl was used as template in a qPCR reaction. Because initially it was not known whether Zbtb11 depletion affects the expression of housekeeping genes, gene expression in Fig. 3b was normalised to the RNA spike-in mix using primers for ERCC 002 and ERCC 003. Following the whole transcriptome analyses at 48 and 72 hours post-*Zbtb11* KO induction, four suitable internal controls were identified (*Hectd1*, *Actb*, *Sel1*, *March7*) that consistently remained unchanged between replicates and data sets, and which were subsequently used for normalisation in Fig. 3c and 5F.

### Immunoblotting

Whole cell lysates were prepared by heating 1 million cells at 95°C in 100μl 1x Laemmli Buffer for 5 minutes. For immunoblots including Mtco1, lysates were prepared in 1x Laemmli Buffer supplemented with Complete protease inhibitors (Roche) by heating at 37°C for 5 minutes, sonicating 30 seconds in a Bioruptor Pico (Diagenode), followed by 5 more minutes of heating at 37°C. Following separation by SDS-PAGE, proteins were transferred onto PVDF membranes, which were subsequently blocked with 5% skimmed milk in [TBS, 0.1% Tween-20]. Antibodies used were: FLAG M2 (F3165, Sigma), Patz1 (sc-292109, Santa Cruz), Lamin B (sc-6216, Santa Cruz), Mrpl48 (14677-1-AP, Proteintech), Ndufc2 (15573-1-AP, Proteintech), Ndufb11 (ab183716, Abcam), Ndufa9 (ab14713, Abcam), Patz1 (sc-292109, Santa Cruz). Mtco1, Ndufb8, Sdhb, Uqcrc2, and Atp5a are part of an OXPHOS antibody cocktail (ab110413, Abcam). Because Mtco1 detection is not as efficient as the rest of the proteins detected with the antibody cocktail (Jha et al. 2016), the membranes were incubated with additional Mtco1 antibody (ab14705, Abcam). Secondary antibodies were HRP-coupled: anti-mouse (P026002-2, Dako), anti-rabbit (A16029, Thermo Fisher Scientific), anti-goat (SAB3700285, Sigma). Chemiluminescence substrates used were WesternSure ECL (926-80100, LI-COR) and SuperSignal West Femto (34094, Fisher Scientific).

### Apoptosis measurement

For intracellular staining of cleaved Caspase-3, cells were fixed in 2% formaldehyde, permeabilised in 90% Methanol, and stored at -20°C. Following rehydration and blocking in PBS containing 0.5% (w/v) BSA, cells were stained for 30 minutes at room temperature with Alexa Fluor 647-conjugated anti-cleaved Caspase-3 (9602, Cell Signaling Technology) and analysed by flow cytometry. The positive control sample was a 1:1 mixture of cells maintained under normal culture conditions and cells heat-shocked at 45°C for 30 minutes, followed by 3 hour recovery at 37°C. For detection of phosphatidylserine externalisation, cells were stained with CF647-coupled Annexin V (4300-0325, Millipore) without fixation, at 37°C.

### ChIP

Cells were fixed in culture media containing 1% formaldehyde (Sigma) for 10 minutes at room temperature. Following fixation, cells were resuspended in [10 mM HEPES pH 7.5,10 mM EDTA, 0.5 mM EGTA, 0.75% Triton X-100, Complete protease inhibitors] and incubated mixing end over end for 10 minutes at 4°C. Nuclei were pelleted by centrifugation, then resuspended and incubated as before in [10 mM HEPES pH 7.5, 200 mM NaCl, 1 mM EDTA, 0.5 mM EGTA, Complete protease inhibtors]. Subsequently nuclei were resuspended in sonication buffer [150 mM NaCl, 25 mM Tris pH 7.5, 5 mM EDTA, 1% Triton X-100, 0.1% SDS, 0.5% Sodium Deoxycholate, Complete protease inhibitors (Roche)], incubated for 30 minutes on ice, and sonicated in a Bioruptor Pico (Diagenode) for 10 cycles (30 seconds on, 30 seconds off). Insoluble chromatin was pelleted by high-speed centrifugation and the supernatant was incubated with the ChIP antibody over night mixing end over end at 4°C. 1μg antibody was used per million sonicated ESCs. Antibodies used were: FLAG M2 (F3165, Sigma) and Zbtb11 (NBP1-80327, Bio-Techne). Following over night incubation, antibody/chromatin complexes were pulled down by mixing 3 hours at 4°C with protein G Dynabeads (Thermo Fihser Scientific) using 5μl beads per μg antibody. Beads were washed 3 times and the DNA was eluted by shaking the beads over night at 65°C in [1xTE, 1% SDS, 0.5mg/ml proteinase K (Sigma)]. Eluted DNA was isolated by phenol/chloroform extraction and ethanol precipitation. Purified DNA was quantified using the Quant-iT PicoGreen kit (Thermo Fisher Scientific).

### ChIP-reChIP

Sonicated chromatin lysates were obtained as described above (see Methods: ChIP) and the protein content was measured using the Bio-Rad RC DC protein measurement kit (cat. no. 5000121). 500μg chromatin was mixed with 5μg anti-Zbtb11 antibody over night. Antibody/chromatin complexes were pulled down with 25μl protein G Dynabeads by mixing for 3 hours at 4°C. The beads were washed three times and the immunoprecipitated chromatin was eluted with 50μl [1xTE, 10mM DTT] by shaking 30 minutes at 37°C. Eluted chromatin was diluted 1:20 with [150 mM NaCl, 25 mM Tris pH 7.5, 5 mM EDTA, 1% Triton X-100, 0.5% Sodium Deoxycholate, Complete protease inhibitors (Roche)], and mixed over night with with [8μg anti-GABPa antibody (Santa Cruz, sc-28312 X) pre-captured on 100μl sheep anti-mouse IgG Dynabeads (by mixing 3 hours at 4°C)]. Beads were washed three times and DNA was eluted over night with 230μl [1xTE, 1% SDS, 1mg/ml proteinase K, 0.3mg/ml RNase A] shaking at 65°C. DNA was extracted using a Qiagen PCR purification kit, and 1/40 was used in each qPCR reaction.

### ChIP-seq

Libraries were prepared from 10ng ChIP DNA using the NEBNext Ultra II DNA Library Prep kit for Illumina (E7645, New England Biolabs). 3 biological replicates were obtained for FLAG-Zbtb11 ChIP-seq in *Zbtb11*^FLAG/FLAG^ E14 cells, and 4 biological replicates for Zbtb11 ChIP-seq in *Zbtb11*^wt^ E14 cells. 50 bases single read sequencing was performed on the Illumina HiSeq 2500 system, obtaining a minimum of 30 million reads for each library. Reads were trimmed and filtered using Cutadapt (Martin 2011) applying a Phred quality threshold of 25 and a minimal read length of 25 bases. Filtered reads were aligned to the mouse genome (NCBI37/mm9 assembly) with Bowtie 1.1.2 (Langmead et al. 2009) keeping only uniquely aligned reads while allowing a maximum of 2 mismatches. MACS2 (Zhang et al. 2008) was set to discard PCR/optical duplicates and was used as peak caller within the Irreproducibility Discovery Rate (IDR) pipeline (Li et al. 2011) implemented by the ENCODE consortium. The final set of peaks was considered the conservative set for IDR <0.02. From this set we also removed blacklisted sites (Carroll et al. 2014). Overlap with UCSC gene annotations and statistical analyses were carried out in R using the Bioconductor packages GenomicRanges and GenomicFeatures (Lawrence et al. 2013), considering promoter regions as 2kb upstream and 500bp downstream of transcription start sites. Normalised read coverage values for consensus peaks were obtained using the DiffBind Bioconductor package (Stark and Brown, 2011).

### Motif analyses

ChIP-seq peaks were centred around the summit and re-sized to 300 bases. Gene promoters were limited to the region -500 to +500 around the TSS. DNA sequences were obtained using the Bioconductor package BSgenome. *De novo* motif search motif matching were performed using the MEME Suite (Bailey et al. 2009).

### RNA-seq

1μg total RNA was used as starting material for each library. rRNA was depleted using the NEBNext rRNA Depletion Kit (E6310, New England Biolabs) and subsequently directional RNA-seq libraries were prepared using the NEBNext Ultra Directional RNA Library Prep Kit for Illumina (E7420, New England Biolabs). 75 bases paired end sequencing was performed on the Illumina HiSeq 2500 system, obtaining a minimum of 31 million reads for each library. Adaptor sequences and bases with a Phred quality score lower than 25 were trimmed from read ends with TrimGalore (Martin 2011), and polyA tails were removed using PRINSEQ (Schmieder and Edwards 2011). Filtered reads were aligned to the mouse transcriptome with STAR (Dobin et al. 2013) using 2-pass mapping, and gene-level counts were generated with HTSeq (Anders et al. 2015, 201). Differential gene expression (DGE) analyses were performed using the Bioconductor package edgeR (generalised linear model approach, quasi-likelihood F-test) (Robinson et al. 2010).

#### DGE analysis

Libraries were prepared from three biological replicates for each of the three sample groups. When all three replicates were used to compare 4OHT- to EtOH-treated *Zbtb11*^lox/lox^ *R26*^ERt2-Cre^ cells, 110 genes were identified as differentially expressed (DE) (FDR < 0.05), 103 of which were down-regulated. Multi-dimensional scaling (MDS) analysis revealed that 2 of the 3 biological replicates were more similar to one another than the remaining third replicate (Supplementary Fig.7a). The fact that this was observed for all sample groups in the same replicate, suggested these differences may be underpinned by technical variability. Removing the distant replicate decreased the dispersion across the entire gene expression range and the common biological coefficient of variation (BCV) value dropped from 17.2% to 6.7% (Supplementary Fig.7b). Repeating the DGE analysis without the distant replicate, 109 of the 110 DE genes found with the 3 replicates were detected, plus an additional 45 DE genes, 43 of which were down-regulated (FDR < 0.05). The effect size at these additional DE genes was smaller (median absolute fold-change 1.84 vs 2.92, p = 1.063e-13, Supplementary Fig. 7c), which is why they were only detected when the BCV is reduced. However, 44 of these genes were found to have very strong Zbtb11 binding sites at their promoters with enrichment values indistinguishable from the binding sites found at the promoters of the DE genes originally detected (p-value = 0.974, see Supplementary Fig. 7d). This suggested that the additional DE genes identified were indeed genuine Zbtb11 targets. Moreover, functional annotation term enrichment analysis with either the restricted (110) or extended (154) list of DE genes gave the same result, the cluster of terms related to mitochondria being the only significant result (FDR < 0.05), with similar enrichment scores of 6.28 and 6.12, respectively. This indicated that although the extended list of DE genes contained 40% more genes, the additional ones were associated with the same functional annotation terms, among which terms related to mitochondria are enriched. We therefore considered the total of 154 DE genes to be the final list of deregulated genes 48 hours post-KO induction (Table S1).

### Pathway mapping and functional enrichment analysis

The Database for Annotation, Visualization and Integrated Discovery (DAVID) (Huang et al. 2009a, 2009b) was used to obtain enriched annotation terms in comparison to the complete list of mouse genes (https://david.ncifcrf.gov/). Pathway mapping was carried out using the Reactome database (version 65) (https://reactome.org/).

### TMRE staining

Cells were trypsinised from maintenance cultures and aliquoted in 12-well plates at 0.5 million cells per well in 0.5ml media. A serial dilution of FCCP was prepared in media containing 400nM TMRE (2x), and 0.5ml of these dilutions was added to the cell suspension for a final concentration of 200nM TMRE and 0, 1, 2, 4, and 8μM FCCP. Cells were returned to the incubator for 20 minutes, after which they were pelleted by low speed centrifugation, washed once and resuspended in [PBS, 2% FBS] for FACS analysis.

### Seahorse assay

Seahorse assay plates were coated with 0.2% gelatin and dried over night in the cell culture incubator. 100,000 cells were plated in each well the same day of the assay and allowed to attach for 5 hours in growth media in the presence of 5μM ROCK inhibitor (13624, Cell Signaling Technology). Subsequently growth media was replaced with Seahorse base media (102353-100, Agilent) containing 2mM sodium pyruvate and 10mM glucose, and plates were incubated for an hour at 37°C and atmospheric CO_2_ concentration. The Seahorse MitoStress kit (103015-100, Agilent) was used to prepare dilutions of the mitochondrial inhibitors which had the following final concentrations: 1μM oligomycin, 0.5μM FCCP, 0.5μM rotenone and antimycin A. Oxygen consumption was measured using a Seahorse XF24 Analyzer with the following program: mix 4 minutes, delay 2 minutes, measure 3 minutes.

#### Normalisation

Control and *Zbtb11* KO cells from each biological replicate were always assayed on the same plate. For normalisation between wells on the same plate, the amount of gDNA was used as a proxy for cell number. At the end of the Seahorse assay, media was replaced with 200μl cell lysis buffer (25mM Tris pH 7.5, 150mM NaCl, 5mM EDTA, 1% Triton X-100, 0.2% SDS, 0.5% Deoxycholate, 0.25mg/ml proteinase K), plates were sealed with adhesive film and incubated over night at 37°C. Plates were briefly centrifuged, and lysates were homogenised by pipetting, followed by 1:10 dilution with ultra pure water. Diluted lysates were transferred to PCR plates and sealed. Proteinase K was inactivated by incubating the diluted lysates at 95°C for 15 minutes, after which 2μl were used in a qPCR reaction. The mean qPCR signal from two different genomic primer pairs (see Supplemental Information for primer sequences) was used to obtain a size factor for each well.

To average oxygen consumption rates (OCR) from all biological replicates, an additional plate-wide adjustment factor was applied. This was calculated based on the expectation that overall, control cells on different plates should have similar maximal OCR after correcting for cell number.

Following normalisation to gDNA, maximal respiration values were averaged across all control samples, and subsequently used to calculate an adjustment factor relative to the inter-replicate mean.

#### Parameter calculation

Following normalisation, mitochondrial respiration parameters were calculated as defined previously (Brand and Nicholls 2011): non-mitochondrial respiration (respiration detected after complete inhibition of mitochondrial respiration with rotenone and antimycin A); basal respiration (respiration measured before the addition of any mitochondrial poisons, minus non-mitochondrial respiration); leak respiration (respiration measured after inhibition of ATP synthase with oligomycin, minus non-mitochondrial respiration); ATP production (basal respiration minus leak and non-mitochondrial respiration); maximal respiration (the maximal respiration measured after treating cells with a pre-determined optimal FCCP amount (0.5μM), minus non-mitochondrial respiration); spare capacity (maximal respiration minus basal respiration); coupling efficiency (ATP production as fraction of basal respiration). Statistical tests were performed using the R package *stats*.

### Mitochondria Isolation

Mitochondria were isolated as described (Corcelli et al. 2010). 25 million cells were lysed in 1ml hypotonic buffer (5mM MOPS pH 7.25, 180mM KCl, 1mM EDTA, EDTA-free protease inhibitors) using a Dounce homogeniser. Nuclei and debris were removed by centrifugation at 1,000xg, and the supernatant was subsequently centrifuged at 10,000xg to pellet mitochondria. Mitochondria were resuspended in the same hypotonic buffer, and following total protein measurement, 50μg aliquots were pelleted by centrifugation at 10,000xg and snap-frozen.

### Blue Native Polyacrylamide Gel Electrophoresis (BN-PAGE)

50μg mitochondria were solubilised in [10mM bis-Tris pH 7.0, 50mM 6-aminohexanoic acid, 2% digitonin] and incubated 20 minutes on ice. Insoluble material was removed by centrifugation and Comassie Brilliant Blue G-250 was added to a final concentration of 0.6%. Gel and electrophoresis buffers used have been previously described (McKenzie et al. 2007). Native complexes were separated for 30 minutes at 150V using blue cathode buffer (15mM bis-Tris pH 7.0, 50mM tricine, 0.02% (w/v) Comassie Brilliant Blue G-250), after which the blue cathode buffer was replaced with clear cathode buffer (15mM bis-Tris pH 7.0, 50mM tricine) and complexes were further separated for 2.5 hours at 250V. In-gel activity assay was performed as previously described (Jha et al. 2016) - following electrophoresis native gels were washed in ice-cold ultra-pure water and incubated mixing in Complex I substrate solution (2mM Tris-HCl pH 7.4, 0.1mg/ml NADH, 2.5mg/ml Nitrotetrazolium Blue Chloride), while monitoring the development of the purple-coloured bands. When the desired contrast was achieved the reaction was stopped with 10% acetic acid. Following imaging, the gels were fixed in [50% methanol, 10% acetic acid] and stained with 0.1% Comassie Brilliant Blue R-250 (in fixation solution). Gels were destained in [40% methanol, 10% acetic acid]. For immunoblotting, following BN-PAGE, native gels were equilibrated in transfer buffer (48mM Tris, 39mM Glycine, 0.03% SDS, 20% Methanol) for 10 minutes and subsequently transferred onto PVDF membrane for 1.5 hours at 350mA using a wet transfer system. Blocking and detection was performed as described above.

## Acknowledgements

We would like to thank our colleagues Marika Charalambous, Reiner Schulz, Cameron Osborne and Ferdinand von Meyenn for discussions and constructive comments. We would also like to thank staff at Guy’s and St Thomas’ NIHR Biomedical Research Centre Genomics and Flow Cytometry facilities for technical assistance, as well as staff maintaining the Rosalind High Performance Compute Cluster.

## Author Contributions

Conceptualization, V.C.S; Methodology, V.C.S; Investigation, B.C.W., L.B., A.A., and V.C.S; Software, A.H., T.C., and V.C.S.; Supervision R.J.O. and V.C.S; Funding Acquisition V.C.S; Writing, V.C.S

## Supplementary Information

Supplementary Fig. 1

Supplementary Fig. 2

Supplementary Fig. 3

Supplementary Fig. 4

Supplementary Fig. 5

Supplementary Fig. 6

Supplementary Fig. 7

Supplementary Table 1 (Excel file). Differentially expressed genes in 48 hour *Zbtb11* KO cells

Supplementary Table 2. Significant Functional Annotation Clusters associated with genes differentially expressed in 48 hour *Zbtb11* KO cells

Supplementary Table 3. Significantly enriched Reactome pathways associated with differentially expressed genes encoding mitochondrial proteins

pPCR Primers

---

**Supplementary Fig. 1.**
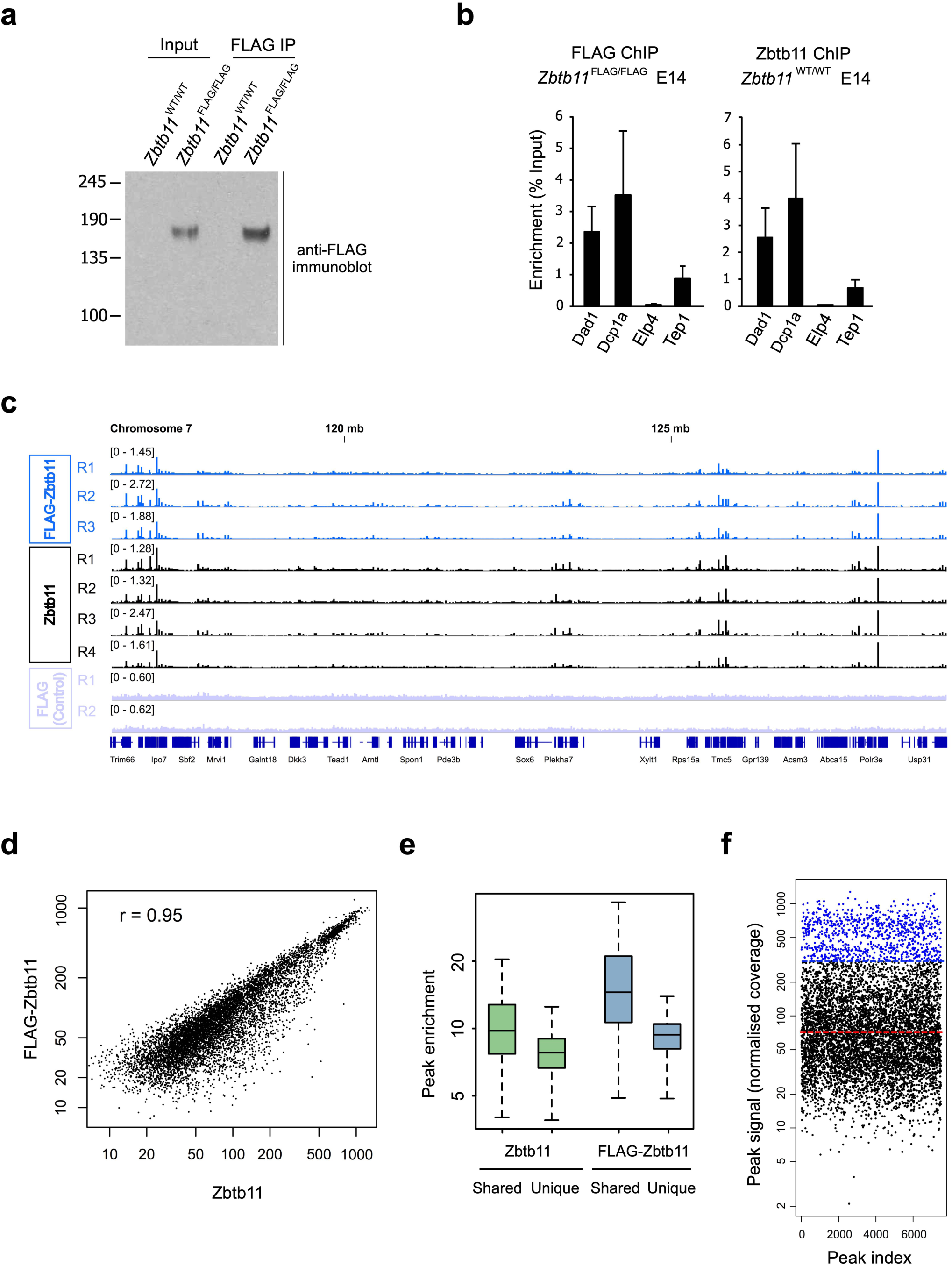
(a) Immunoblot with anti-FLAG antibodies showing specific and efficient immunoprecipitation (IP) of FLAG-Zbtb11 from chromatin lysates (Input) of *Zbtb11*^FLAG/FLAG^ E14 cells but not from lysates of the wild type parental line. (b) qPCR analysis of FLAG ChIP samples from *Zbtb11*^FLAG/FLAG^ E14 cells and of anti-Zbtb11 ChIP from the parental wild type E14 line. Bars show mean and SD of ChIP enrichment values (as percentage of input) obtained from three biological replicates. The panel of primers amplify the promoter regions at the indicated genes. Note the similar distribution of enrichment values obtained with the two different approaches. (c) ChIP-seq signal across the entire chromosome 7 illustrating the reproducible enrichment obtained for both FLAG-Zbtb11 ChIP in *Zbtb11*^FLAG/FLAG^ E14 cells and Zbtb11 ChIP in the parental wild type E14 line. By contrast, no enrichment was generated by the control ChIP experiment using FLAG antibodies in the parental E14 line. (d) Correlation of Zbtb11 and FLAG-Zbtb11 ChIP-seq signal at consensus peaks. Plotted values represent mean normalised coverage obtained by averaging the individual biological replicates from each dataset. Pearson’s r is shown. (e) Box plot showing the distributions of FLAG-Zbtb11 and Zbtb11 ChIP enrichment values (as obtained during peak calling, see Methods) at peaks that are common or unique to the two datasets. Note that peaks detected with only one of the two approaches are significantly weaker than the common peaks, which makes them more likely to be false positives. (f) Distribution of Zbtb11 ChIP signal at consensus peaks. Outlier values (one and a half-time the interquartile range over the third quartile) which we use to designate Zbtb11-high peaks are shown in blue. The median of the distribution is shown by the red line.

**Supplementary Fig. 2.**
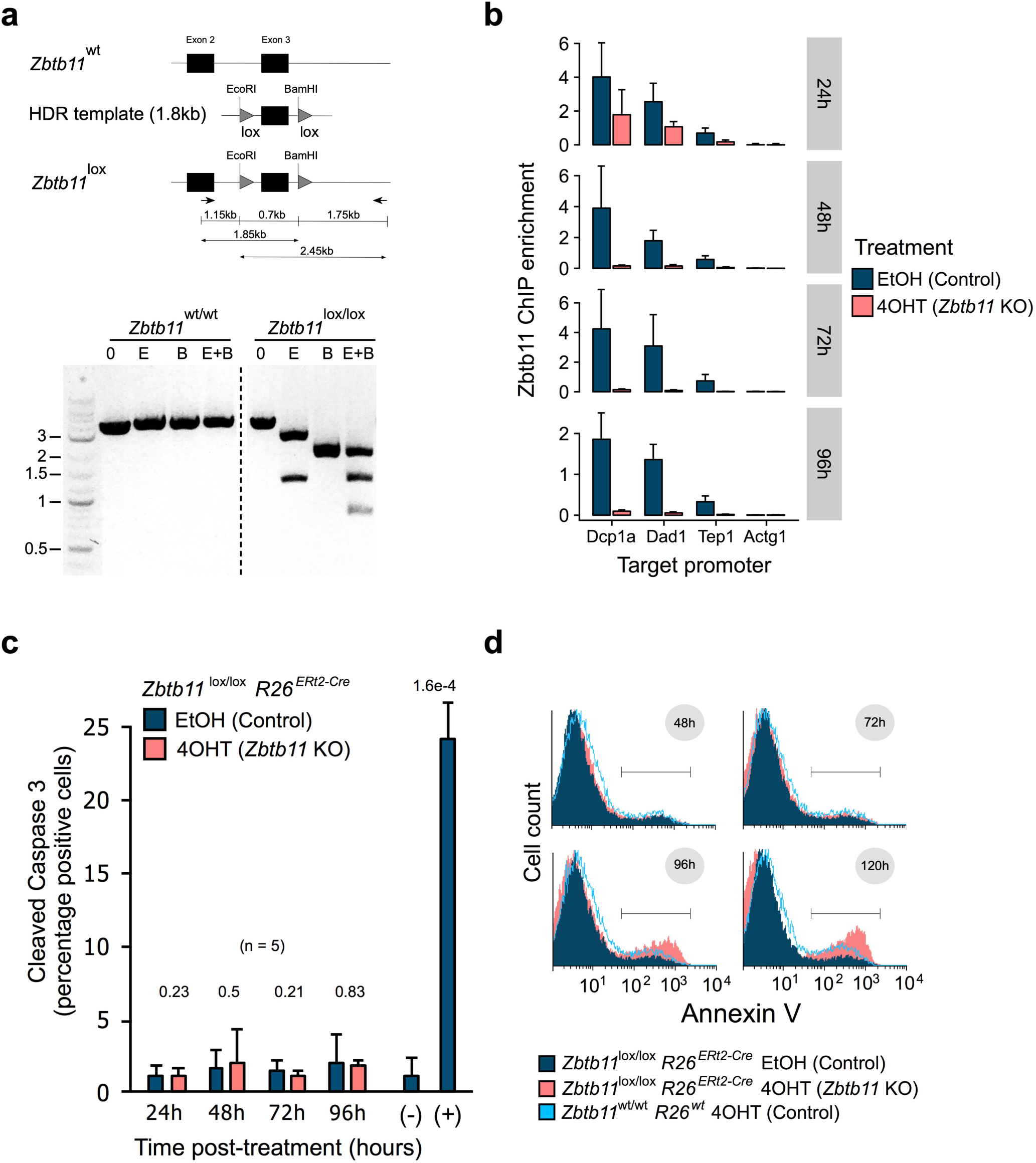
(a) Restriction enzyme digest analysis of the *Zbtb11* locus in *Zbtb11*^lox/lox^ *Rosa26*^ERt2-Cre^ ESC line. Successful insertion of the two loxP sites flanking exon 3 introduces an ectopic EcoRI site in intron 2 and a BamHI site in intron 3. A 3.6kb region spanning intron 2, exon 3 and part of intron 3 was amplified from gDNA using primers outside the region of homology with the donor template (HDR). Lower panel - DNA agarose gel showing that the PCR amplicon from the parental wild type line is not digested by either EcoRI or BamHI (left), while the amplicons from the recombinant homozygous line are digested by both EcoRI and BamHI (right). The dotted vertical line in the lower panel indicates where intervening lanes have been removed from the image. (b) ChIP-qPCR with anti-Zbtb11 antibodies in *Zbtb11*^lox/lox^ *Rosa26*^ERt2-Cre^ cells treated with either EtOH (control) or 4OHT (*Zbtb11* KO) (treatment duration indicated on the right). The primers amplify promoter regions at the indicated genes. Bars show mean and SD of three biological replicates. Note the complete depletion of Zbtb11 protein 48 hours after KO is induced. (c) Quantification of apoptotic cells following Zbtb11 KO. *Zbtb11*^lox/lox^ *Rosa26*^ERt2-Cre^ cells were treated with either EtOH or 4OHT and samples were subsequently collected and fixed at 24 hour intervals. Cells were then stained with fluorescently labelled antibodies to Caspase-3 and analysed by flow cytometry to measure the fraction of Caspase-3 positive cells. Bars show mean and SD of 5 biological replicates, with two-tailed t-test p-values above. Note that this approach did not reveal an increase in apoptotic cells despite being able to specifically and efficiently detect apoptosis in a positive control sample made up of cells submitted to heat-shock (+). (d) Representative histograms showing the distribution of Annexin V-binding cells as measured by flow cytometry, which was used as a marker of cell death. Line interval demarcates Annexin V-positive cells. Note that an increase in Annexin V-binding cells becomes evident 96 hours after *Zbtb11* KO induction.

**Supplementary Fig. 3.**
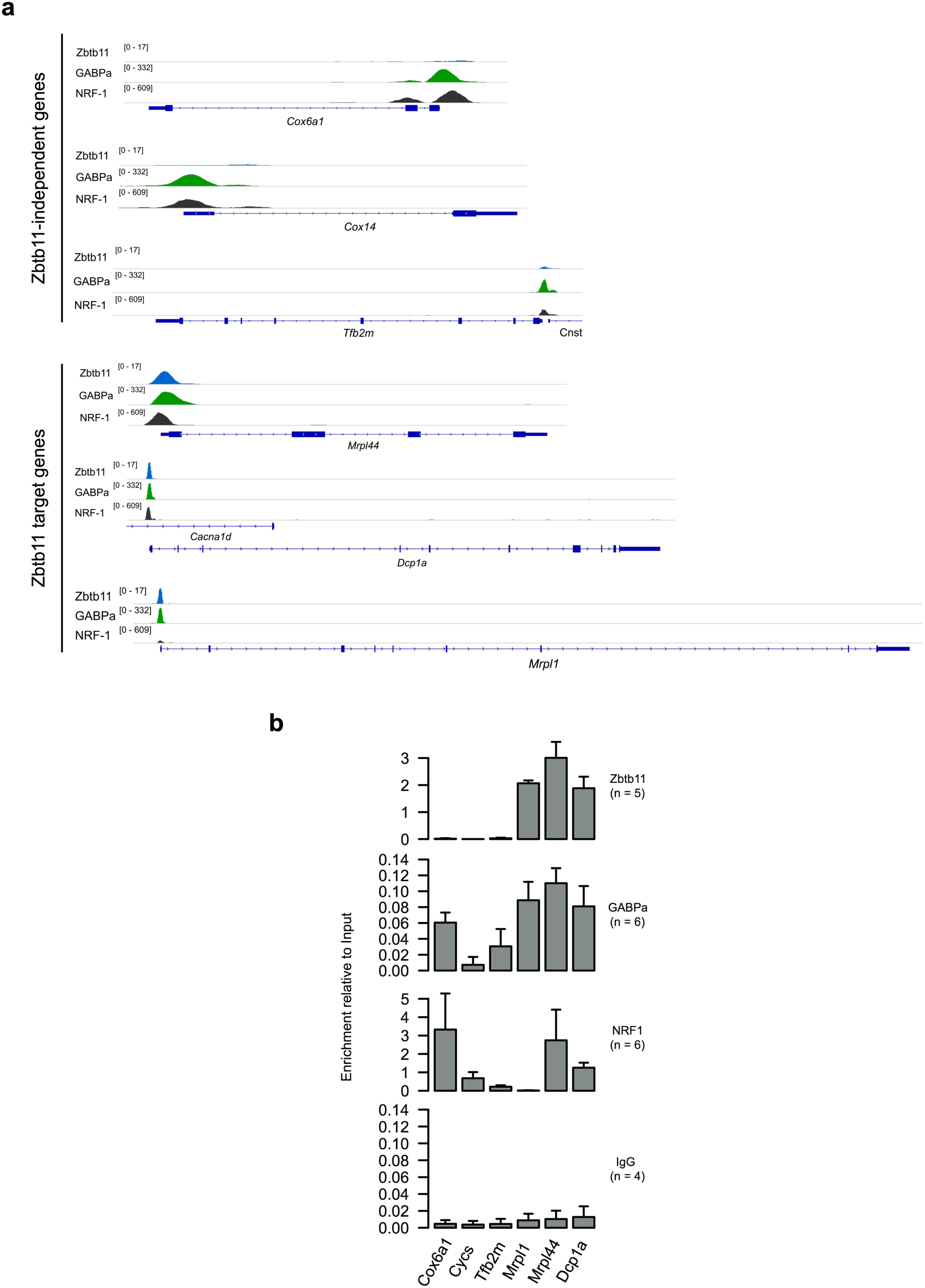
(a) Read coverage tracks for Zbtb11, GABPa and NRF-1 ChIP-seq in WT mouse ESCs. Three Zbtb11-independent loci (top) and three Zbtb11-dependent loci (bottom) are shown. (b) ChIP-qPCR for Zbtb11, GABPa, NRF-1 and control rabbit IgG. Bars are mean and SD of enrichment values relative to Input detected with primers amplifying the promoter regions of the genes indicated underneath.

**Supplementary Fig. 4.**
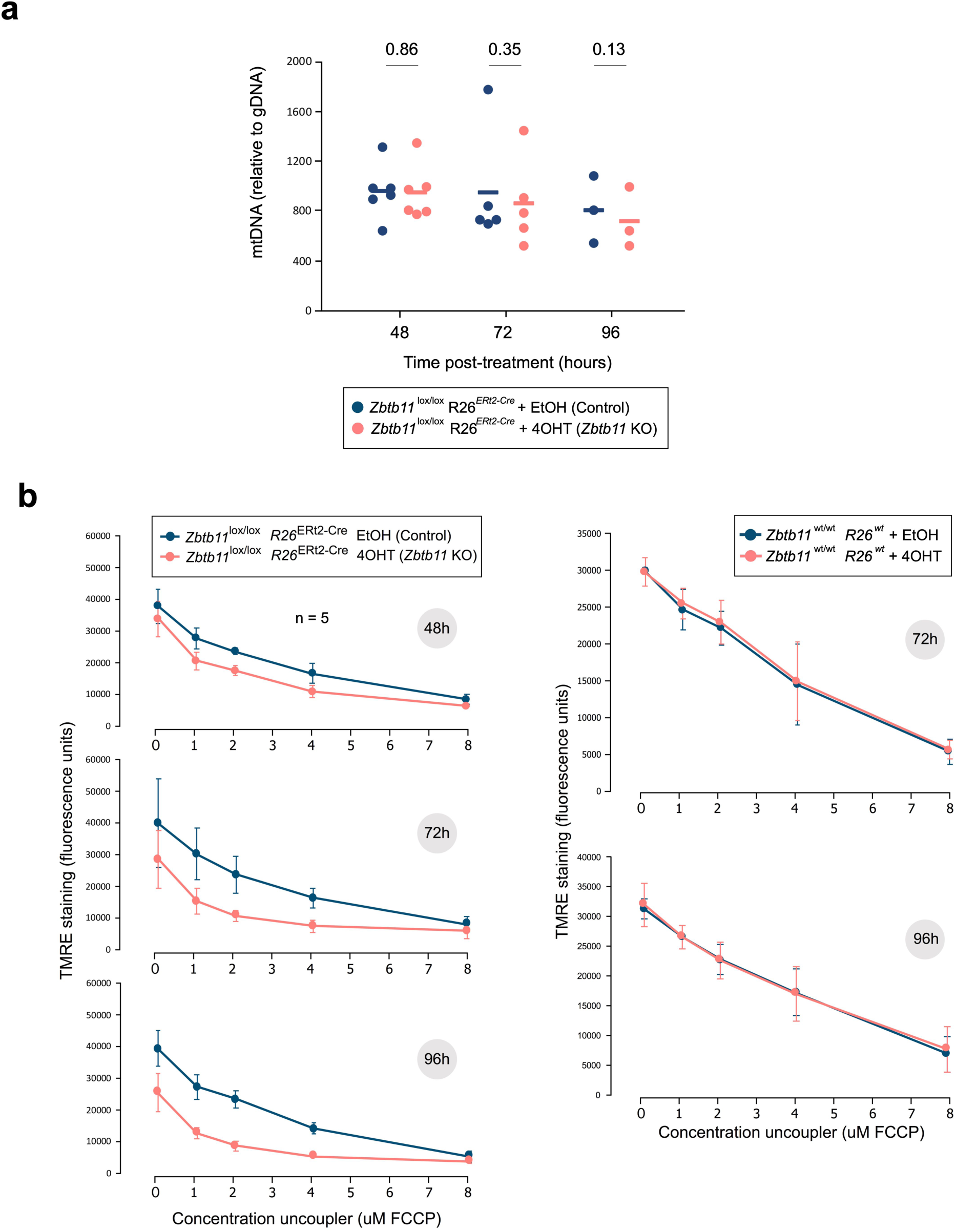
(a) qPCR quantification of mtDNA content relative to gDNA following *Zbtb11* KO induction. (b) Increased sensitivity of the MMP to mitochondrial uncoupler treatment in *Zbtb11* KO cells. Panels to the left - control and *Zbtb11* KO cells (time post-4OHT treatment indicated on the right) were stained with TMRE in the presence of increasing amounts of FCCP, and analysed by flow cytometry. Plots show TMRE mean fluorescence intensity against FCCP concentration (mean ± SD, n = 4-7 biological replicates). Panels to the right - to control for potential non-specific effects of the 4OHT treatment on the MMP, the TMRE staining and FCCP titration was carried out on EtOH- and 4OHT-treated *Zbtb11*^wt/wt^ *Rosa26^wt^* cells, showing that 4OHT treatment of wild type cells does not affect the MMP.

**Supplementary Fig. 5.**
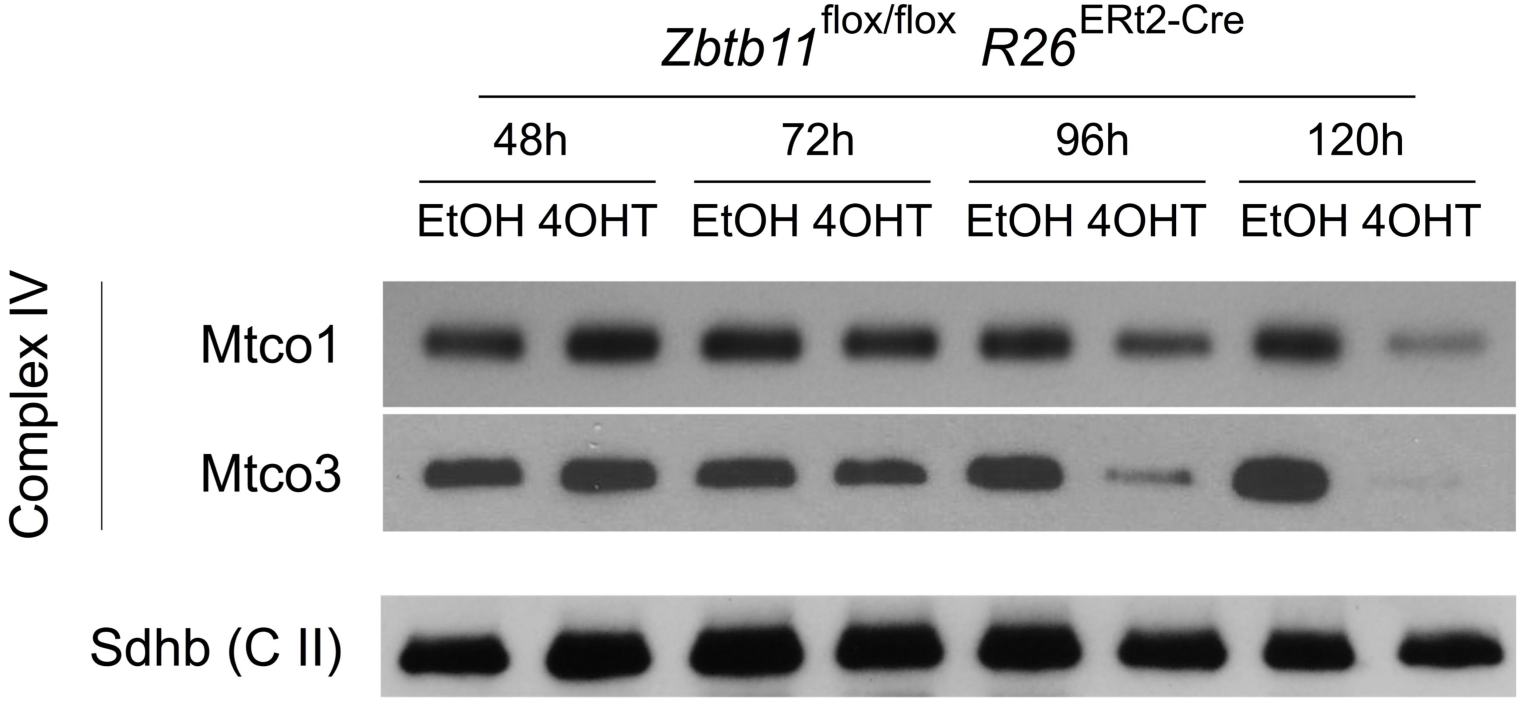
SDS-PAGE followed by immunoblotting of mitochondrial extracts from EtOH-treated (control) and 4OHT-treated (*Zbtb11* KO) *Zbtb11*^lox/lox^ *Rosa26*^ERt2-Cre^ cells, showing the mitochondria-encoded proteins Mtco1 and Mtco3 are down-regulated from 96 hours post-KO induction onwards. The nuclear-encoded complex II subunit Sdhb is shown as loading reference.

**Supplementary Fig. 6.**
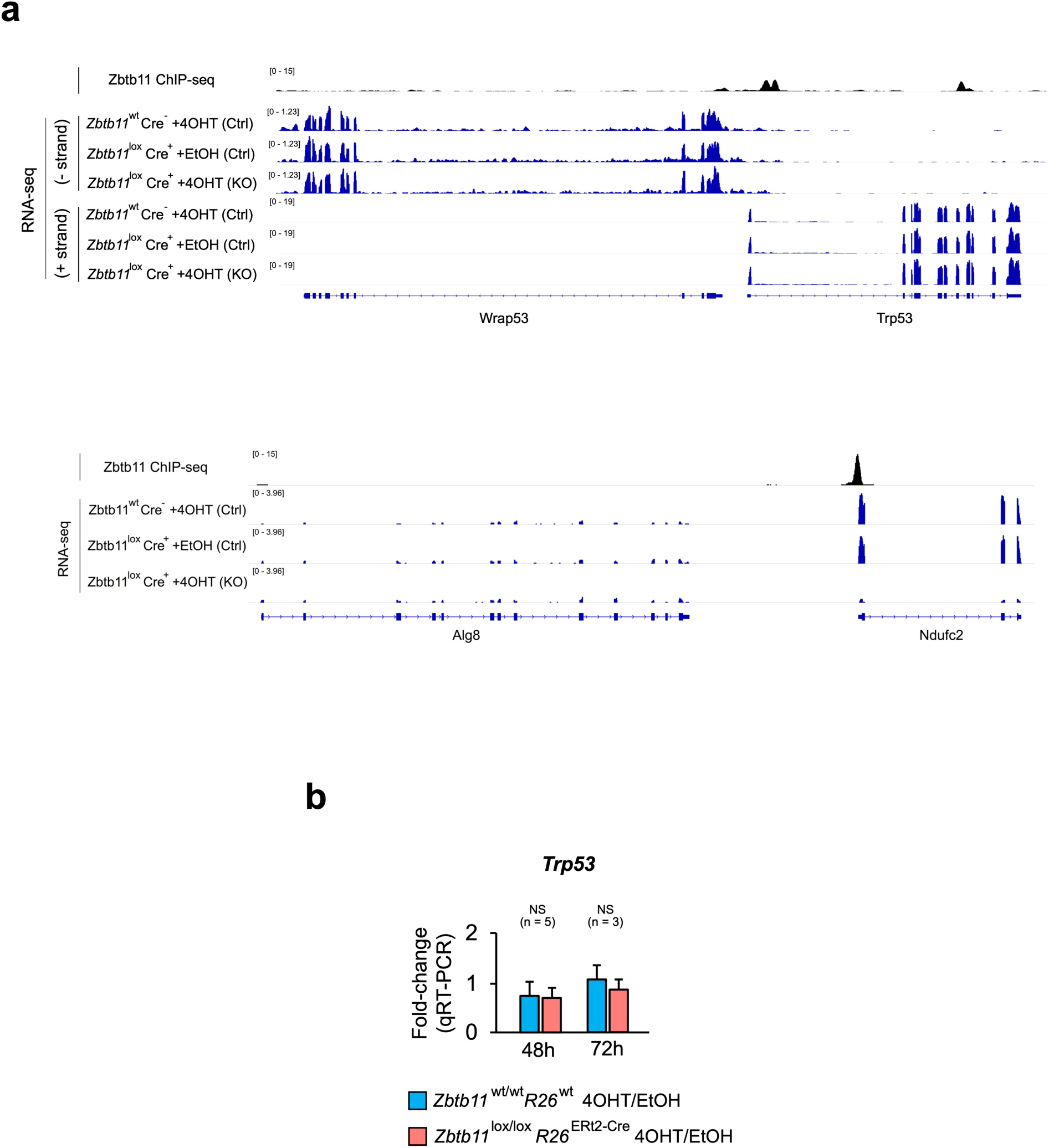
(a) Visualisation of normalised read coverage tracks for Zbtb11 ChIP-seq in control cells, and for RNA-seq in control and *Zbtb11* KO cells. Top panel – locus encoding p53 (*Trp53*) showing no transcriptional changes. Bottom panel - *Ndufc2* locus shown for comparison, illustrating transcriptional down-regulation. (b) qRT-PCR analysis of *Trp53* transcription in control and *Zbtb11* KO cells, 48 and 72 hours post-KO induction. Bars show mean and SD of 3-5 biological replicates.

**Supplementary Fig. 7.**
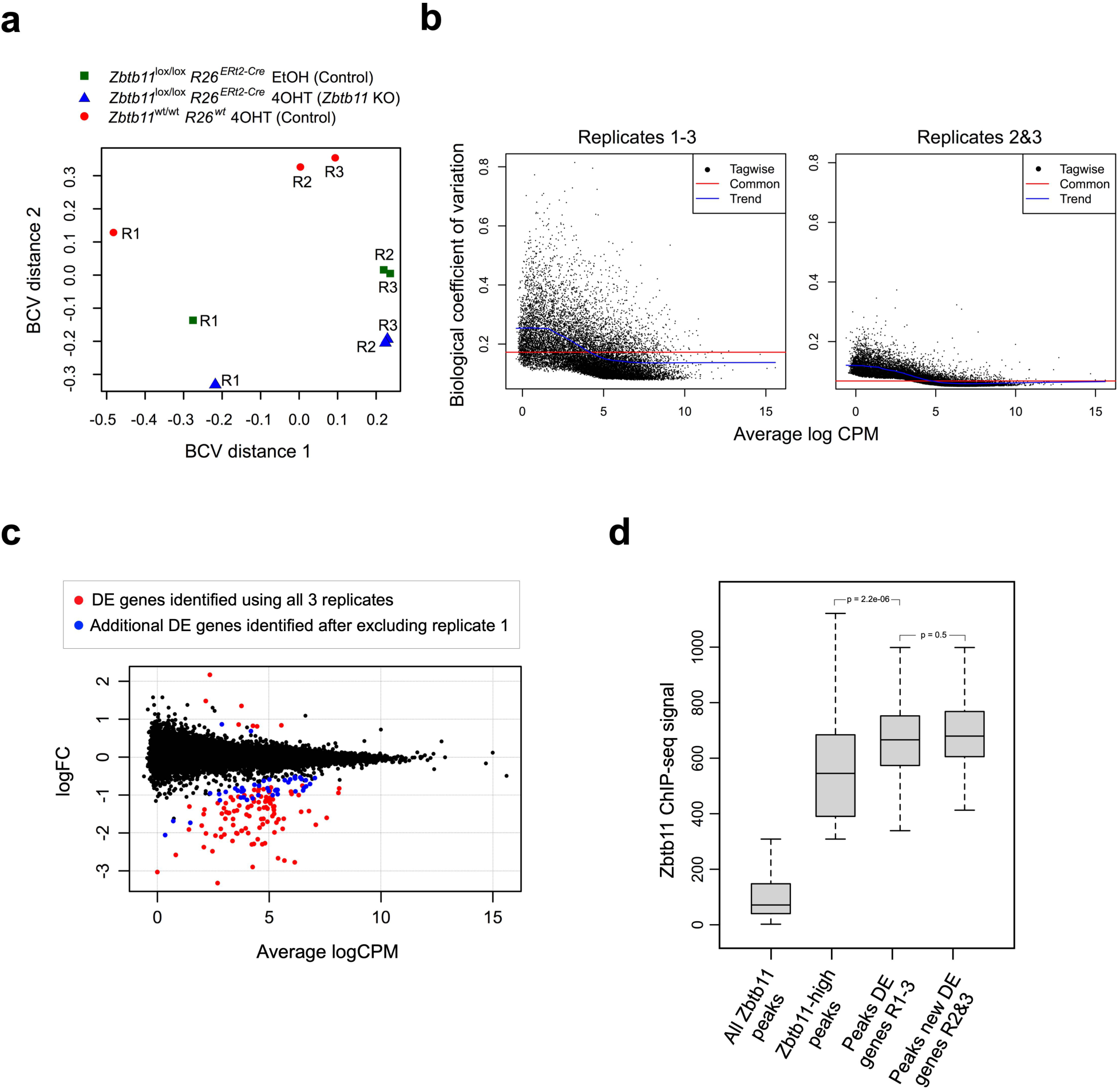
(a) Multidimensional scaling (MDS) plot of the RNA-seq libraries. (b) Biological coefficient of variation (BCV) plotted against transcript abundance (mean counts per million) when all three RNA-seq replicates are included in the analysis (left) or when replicate 1 is removed (right). (c) Plot of log2-transformed fold-change against transcript abundance obtained after removing RNA-seq replicate 1 from the differential gene expression analysis. In red are shown DE genes found by using all three replicates, while additional DE genes identified after removal of replicate 1 are shown in blue. (d) Boxplot of Zbtb11 ChIP-seq signal showing that peaks at promoters of the additional DE genes identified once RNA-seq replicate 1 is removed are as strong as the peaks found at the DE genes initially identified with all three RNA-seq replicates. P-values for pair-wise comparisons of the distributions are shown above the plot and were obtained by performing a Wilcoxon rank sum test.

**Supplementary Table 2.** Significant Functional Annotation Clusters associated with genes differentially expressed in 48 hour *Zbtb11* KO cells.

| Annotation Cluster 1 |  | Enrichment Score: 6.28 |  |  |  |
| --- | --- | --- | --- | --- | --- |
| Category | Term | Count | % | PValue | FDR |
| UP_KEYWORDS | Transit peptide | 16 | 14.95 | 1.54E-08 | 1.80E-05 |
| UP_KEYWORDS | Mitochondrion | 21 | 19.63 | 8.52E-08 | 9.97E-05 |
| GOTERM_CC_DIRECT | Mitochondrion | 24 | 22.43 | 3.02E-06 | 0.0035 |

| Annotation Cluster 2 |  | Enrichment Score: 2.14 |  |  |  |
| --- | --- | --- | --- | --- | --- |
| Category | Term | Count | % | PValue | FDR |
| REACTOME_PATHWAY | Mitochondrial translation elongation | 6 | 5.61 | 4.99E-05 | 0.056 |
| REACTOME_PATHWAY | Mitochondrial translation termination | 6 | 5.61 | 5.60E-05 | 0.063 |
| GOTERM_CC_DIRECT | Intracellular ribonucleoprotein complex | 8 | 7.48 | 7.61E-04 | 0.880 |
| UP_KEYWORDS | Ribosomal protein | 6 | 5.61 | 0.003 | 3.080 |
| GOTERM_CC_DIRECT | Ribosome | 5 | 4.67 | 0.012 | 12.851 |
| UP_KEYWORDS | Ribonucleoprotein | 6 | 5.61 | 0.015 | 16.107 |
| GOTERM_MF_DIRECT | Poly(A) RNA binding | 10 | 9.35 | 0.077 | 61.867 |
| KEGG_PATHWAY | Ribosome | 3 | 2.80 | 0.127 | 70.210 |
| GOTERM_MF_DIRECT | Structural constituent of ribosome | 4 | 3.74 | 0.128 | 80.709 |
| GOTERM_BP_DIRECT | Translation | 4 | 3.74 | 0.286 | 98.840 |

| Annotation Cluster 3 |  | Enrichment Score: 1.90 |  |  |  |
| --- | --- | --- | --- | --- | --- |
| Category | Term | Count | % | PValue | FDR |
| GOTERM_BP_DIRECT | Methylation | 5 | 4.67 | 0.008 | 10.041 |
| GOTERM_MF_DIRECT | Methyltransferase activity | 5 | 4.67 | 0.008 | 9.332 |
| UP_KEYWORDS | Methyltransferase | 5 | 4.67 | 0.009 | 10.497 |
| UP_KEYWORDS | S-adenosyl-L-methionine | 4 | 3.74 | 0.038 | 36.486 |

**Supplementary Table 3.** Significantly enriched Reactome pathways associated with differentially expressed genes encoding mitochondrial proteins.

| Pathway identifier | Pathway name | Found | Total | p-Value | FDR | Reactions | Found |
| --- | --- | --- | --- | --- | --- | --- | --- |
| R-HSA-5419276 | Mitochondrial translation termination | 8 | 88 | 9.73E-10 | 3.21E-08 | 5 | Mrpl3<br>Mrpl44<br>Mrpl1<br>Mrpl13<br>Mrpl58<br>Mrpl48<br>Mrps31<br>Mrpl30 |
| R-HSA-5368286 | Mitochondrial translation initiation | 8 | 88 | 9.73E-10 | 3.21E-08 | 3 | Mrpl3<br>Mrpl44<br>Mrpl1<br>Mrpl13<br>Mrpl58<br>Mrpl48<br>Mrps31<br>Mrpl30 |
| R-HSA-5389840 | Mitochondrial translation elongation | 8 | 88 | 9.73E-10 | 3.21E-08 | 5 | Mrpl3<br>Mrpl44<br>Mrpl1<br>Mrpl13<br>Mrpl58<br>Mrpl48<br>Mrps31<br>Mrpl30 |
| R-HSA-5368287 | Mitochondrial translation | 8 | 94 | 1.62E-09 | 4.06E-08 | 13 | Mrpl3<br>Mrpl44<br>Mrpl1<br>Mrpl13<br>Mrpl58<br>Mrpl48<br>Mrps31<br>Mrpl30 |
| R-HSA-72766 | Translation | 9 | 294 | 8.47E-07 | 1.69E-05 | 14 | Mrpl3<br>Mrpl44<br>Mrpl1<br>Mrpl13<br>Mrpl58<br>Mrpl48<br>Mrps31<br>Tars2<br>Mrpl30 |
| R-HSA-611105 | Respiratory electron transport | 5 | 101 | 3.18E-05 | 5.09E-04 | 13 | Ndufa12<br>Taco1<br>Ndufc2<br>Ndufs7<br>Ndufaf1 |
| R-HSA-6799198 | Complex I biogenesis | 4 | 55 | 4.82E-05 | 6.74E-04 | 11 | Ndufa12<br>Ndufc2<br>Ndufs7<br>Ndufaf1 |
| R-HSA-163200 | Respiratory electron transport, ATP synthesis by chemiosmotic coupling, and heat production by uncoupling proteins. | 5 | 124 | 8.35E-05 | 0.001 | 13 | Ndufa12<br>Taco1<br>Ndufc2<br>Ndufs7<br>Ndufaf1 |
| R-HSA-1428517 | The citric acid (TCA) cycle and respiratory electron transport | 5 | 175 | 4.09E-04 | 0.00449 | 13 | Ndufa12<br>Taco1<br>Ndufc2<br>Ndufs7<br>Ndufaf1 |

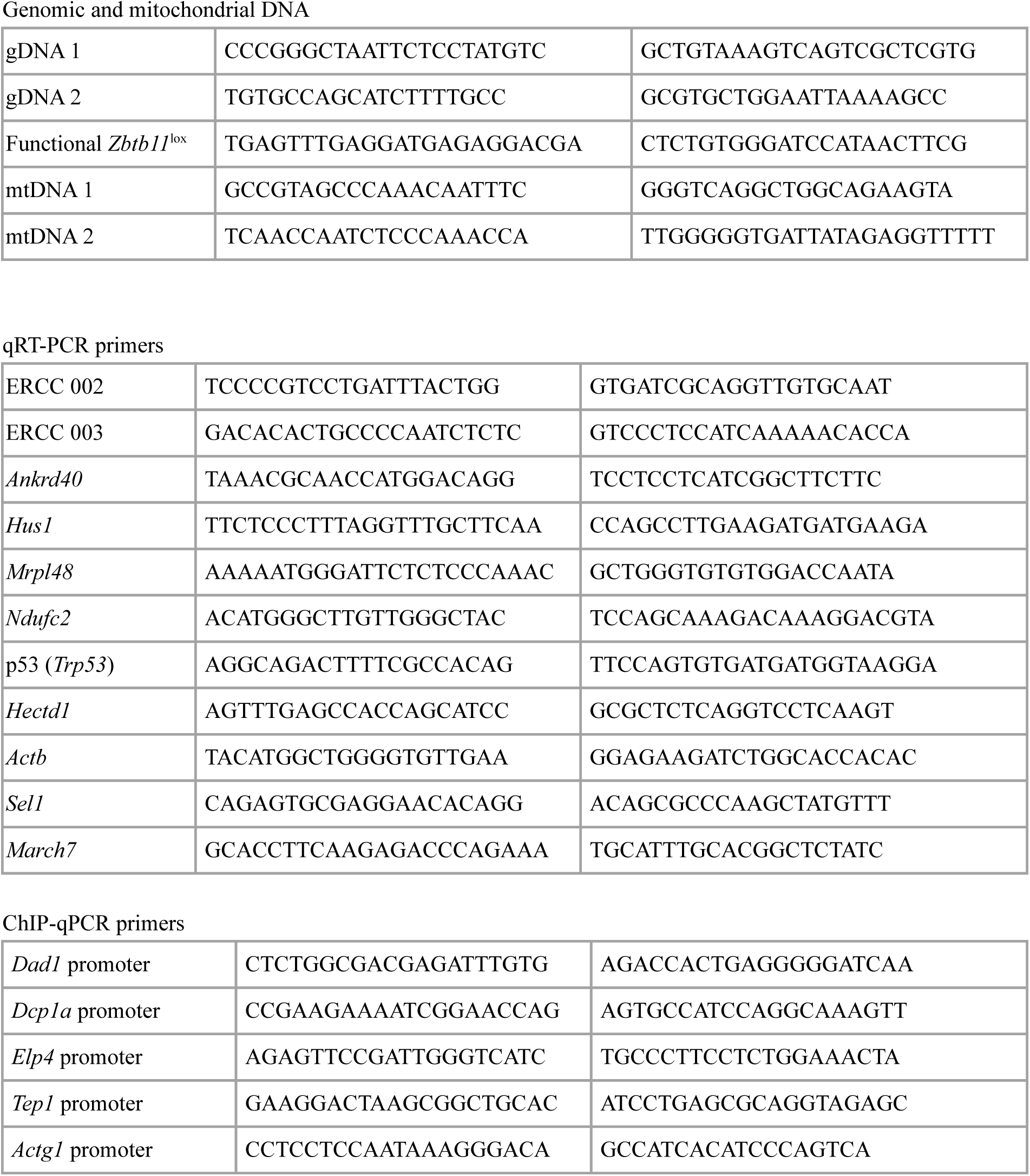
qPCR Primers.

